# Dysregulated alveolar epithelial cell progenitor function and identity in Hermansky-Pudlak syndrome

**DOI:** 10.1101/2023.06.17.545390

**Authors:** Joanna Y. Wang, Sylvia N. Michki, Sneha Sitaraman, Brandon J. Banaschewski, Reshma Jamal, Jason J. Gokey, Susan M. Lin, Jeremy B. Katzen, Maria C. Basil, Edward Cantu, Jonathan A. Kropski, Jarod A. Zepp, David B. Frank, Lisa R. Young

**Affiliations:** Division of Pulmonary, Allergy, Critical Care, and Sleep Medicine, Department of Medicine, University of Pittsburgh School of Medicine, Pittsburgh, Pennsylvania; Division of Pulmonary, Allergy, and Critical Care, Department of Medicine, Perelman School of Medicine, University of Pennsylvania, Philadelphia, Pennsylvania; Division of Cardiology, Department of Pediatrics, Children’s Hospital of Philadelphia, Philadelphia, Pennsylvania, and; Division of Pulmonary and Sleep Medicine, Department of Pediatrics, Children’s Hospital of Philadelphia, Philadelphia, Pennsylvania; Division of Allergy, Pulmonary, and Critical Care Medicine, Department of Medicine, Vanderbilt University, Nashville, Tennessee; Lung Biology Institute, University of Pennsylvania, Philadelphia, Pennsylvania; Penn Cardiovascular Institute, University of Pennsylvania, Philadelphia, Pennsylvania; Department of Surgery, Perelman School of Medicine, University of Pennsylvania, Philadelphia, Pennsylvania

## Abstract

Hermansky-Pudlak syndrome (HPS) is a genetic disorder of endosomal protein trafficking associated with pulmonary fibrosis in specific subtypes, including HPS-1 and HPS-2. Single mutant HPS1 and HPS2 mice display increased fibrotic sensitivity while double mutant HPS1/2 mice exhibit spontaneous fibrosis with aging, which has been attributed to HPS mutations in alveolar epithelial type II (AT2) cells. We utilized HPS mouse models and human lung tissue to investigate mechanisms of AT2 cell dysfunction driving fibrotic remodeling in HPS. Starting at 8 weeks of age, HPS mice exhibited progressive loss of AT2 cell numbers. HPS AT2 cell function was impaired *ex vivo* and *in vivo*. Incorporating AT2 cell lineage tracing in HPS mice, we observed aberrant differentiation with increased AT2-derived alveolar epithelial type I cells. Transcriptomic analysis of HPS AT2 cells revealed elevated expression of genes associated with aberrant differentiation and p53 activation. Lineage tracing and organoid modeling studies demonstrated that HPS AT2 cells were primed to persist in a Krt8^+^ reprogrammed transitional state, mediated by p53 activity. Intrinsic AT2 progenitor cell dysfunction and p53 pathway dysregulation are novel mechanisms of disease in HPS-related pulmonary fibrosis, with the potential for early targeted intervention before the onset of fibrotic lung disease.

## Introduction

Hermansky-Pudlak syndrome (HPS) is an autosomal recessive disorder of endosomal protein trafficking. At birth, patients with HPS do not exhibit pulmonary disease, but progressive pulmonary fibrosis typically develops in young adulthood in HPS-1 and HPS-4, and in childhood in HPS-2 (1). Notably, pulmonary fibrosis has not been reported in HPS-3, which is the second most common HPS subtype after HPS1. We originally demonstrated that HPS mutations specifically in the alveolar epithelium underlie fibrotic susceptibility (2). Evidence suggests that dysfunctional AT2 cells, which act as stem cells within the alveolus and produce and store surfactant, drive fibrotic remodeling in other genetic and sporadic forms of pulmonary fibrosis (3–5). In HPS-related pulmonary fibrosis (HPS-PF), studies from several groups have implicated apoptosis, impaired autophagy, and mitochondrial dysfunction in AT2 cells as potential contributing pathways (2, 6–9). Despite these advances, the development of effective, targeted therapies for HPS-PF has been limited by an incomplete understanding of the consequences of lysosomal and endosomal trafficking defects on AT2 cell function, as well as targetable mechanisms underlying HPS AT2 cell dysfunction. In addition, it remains unclear whether there are common final pathways of AT2 cell dysfunction shared across sporadic and genetic fibrotic lung diseases to inform the generalizability of therapies for pulmonary fibrosis that are currently available or in development.

In this study, we examined HPS AT2 cells over time to characterize the cellular and molecular mechanisms that drive AT2 cell dysfunction and their contribution to the fibrotic response. We utilized five murine models of HPS, including the double mutant HPS1/2 murine model, which exhibits spontaneous progressive fibrosis with aging, as well as the single mutant HPS1 and HPS2 models, which best reflect human disease genetics while exhibiting fibrotic sensitivity without spontaneous fibrosis (**Table 1**) (8, 10–12). We report that in HPS genotypes associated with pulmonary fibrosis, impaired AT2 cell progenitor function occurs early, brought about in part by p53 activation. Starting at eight weeks of age in HPS mice, we show dysregulated maintenance of the alveolar epithelium, with progressive loss of AT2 cells in association with impaired proliferation and aberrant proliferation and persistence of a Krt8^+^ reprogrammed transitional cell state. We also demonstrate the presence of senescent and KRT8^+^ epithelial cells in lung tissue from a patient with HPS-1. Our findings reveal a novel therapeutic target in HPS-PF with the potential for early intervention and suggest the existence of convergent mechanisms of disease between HPS-PF and other sporadic and genetic fibrotic lung diseases.

**Table 1.** Murine models of Hermansky-Pudlak syndrome. . Table adapted from Young et al. (2). ^a^ HPS2 transgenic (TG) model, HPS2 with transgenic correction of AP3b1 in lung epithelium (2). Other notable models include HPS3, which exhibits naturally occurring mutations in *Hps3* and is not associated with pulmonary fibrosis in human disease or mouse studies. Definition of abbreviations: AP-3, adaptor protein 3.

| **HPS mouse models** | **HPS1** | **HPS2** | **HPS1/2** |
| --- | --- | --- | --- |
| **Model** | Naturally occurring *Hps1* mutations | Naturally occurring *Ap3b1* mutations | HPS1 mouse x HPS2 mouse |
| **Human disease correlate** | HPS-1 | HPS-2 | Not seen in patients |
| **Rationale for study** | Most common subtype of HPS associated with pulmonary fibrosis | Gene encodes AP-3, protein most amenable to study; allows for comparison to HPS2 TG<sup>a</sup> | Spontaneous fibrotic phenotype |
| **Fibrotic phenotype** | Fibrotic sensitivity to bleomycin | Fibrotic sensitivity to bleomycin | Spontaneous, progressive fibrosis by 48 weeks |

## Results

### Loss of AT2 cells in HPS mice starting at 8 weeks of age

We first confirmed that double mutant HPS1/2 mice developed spontaneous, progressive fibrosis with aging as previously described (**Supplemental Figure 1A-D**) (11–13). We then utilized immunofluorescence (IF) for the AT2 cell marker, proSP-C, and the alveolar epithelial type I (AT1) cell marker, AGER, to examine the alveolar epithelium in aged HPS1/2 mice compared to wild type (WT) at 48 weeks of age. In addition to clusters of AT2 cell hyperplasia, we observed regions of near-total loss of proSP-C^+^ AT2 cells in HPS1/2 mice (**Figure 1A**). To evaluate the time course of AT2 cell loss, we utilized IF for proSP-C to quantify AT2 cells in WT and HPS1/2 mice at 4, 8, 24, and 48 weeks of age (**Figure 1B-D**). Starting at eight weeks of age, we found a significant decrease in the percentage of proSP-C^+^ AT2 cells per total DAPI positive cells in HPS1/2 mice, with 10.3 ± 0.4% in HPS1/2 mice compared to 13.9 ± 0.3% in WT (adjusted *p* = .02) (**Figure 1D**). These percentages corresponded to a mean 31.0 ± 10.2 cells per high powered field (hpf) in HPS1/2 mice and 56.8 ± 10.4 AT2 cells per hpf in WT mice (*p* < .0001). There was progressive loss of AT2 cells with aging in HPS1/2 mice, which exhibited 8.1 ± 0.5% proSP-C^+^ cells at 48 weeks of age (adjusted *p* = .001), as well as WT mice with aging, with 11.5 ± 0.2% at 48 weeks of age (adjusted *p* = .002) (**Figure 1D**).

**Figure 1.**
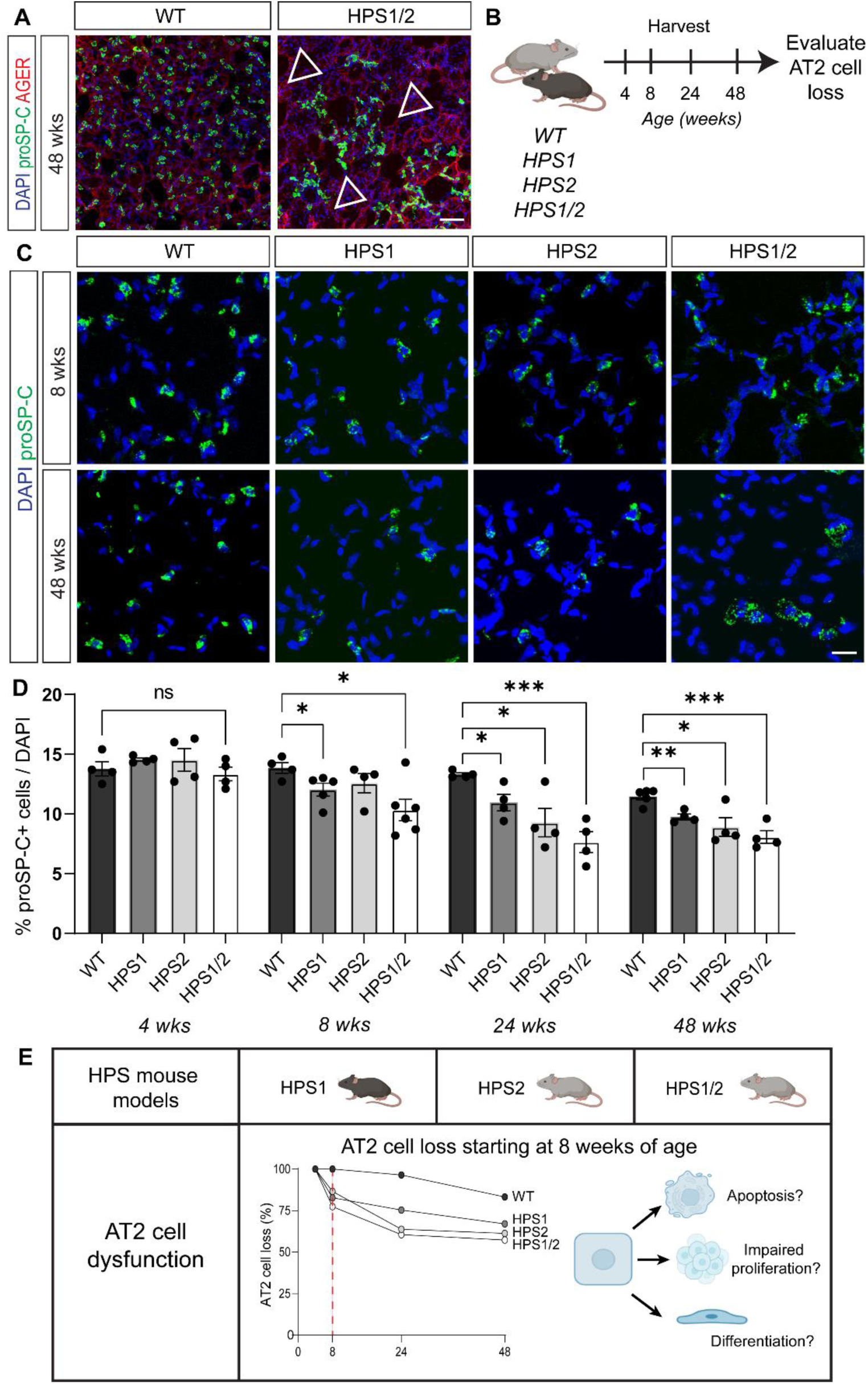
Loss of alveolar type II epithelial (AT2) cells in HPS mice starting at 8 weeks of age. (A) Immunofluorescence (IF) staining of whole-mount lung issue for pro-SPC and AGER in WT and HPS1/2 mice at 48 weeks of age. Arrows indicate regions of AT2 cell loss. (B) Schematic to evaluate AT2 cell loss in WT, HPS1, HPS2, and HPS1/2 mice over time. (C) Staining of paraffin-embedded lung tissue for proSP-C in WT, HPS1, HPS2, and HPS1/2 mice at 8 and 48 weeks of age. (D) Quantification of percentage of proSP-C^+^ cells as a percentage of total cells (by DAPI) in WT, HPS1, HPS2, and HPS1/2 mice at 4, 8, 24, and 48 weeks of age. (E) Schematic of AT2 cell loss in HPS mice and possible etiologies. DAPI stains nuclei (blue). All quantification data are represented as mean ± SEM. Statistics using two-tailed unpaired Student’s *t* tests: ns, not significant; adjusted * *p* < .05; ** *p* < .01, *** *p* < .001 after Benjamini Hochberg correction for multiple comparisons. n = 4-6 per group per time point. Scale bars in (A), 50 μm; (C), 20 μm. Schematics created with Biorender.com.

As HPS1/2 mice contain mutations in both *Hps1* and *Ap3b1* (*Hps2*), we next assessed HPS1 and HPS2 mice to evaluate if the phenotype of AT2 cell loss was dependent on the combinatorial experimental HPS model or also occurred in mice with disruption in a single HPS gene. Significant AT2 cell loss was also seen in single mutant HPS1 (adjusted *p* < .0001) and HPS2 mice (adjusted *p* = .037), respectively (**Figure 1C, D**). The difference in AT2 cell numbers was detected starting at eight weeks of age in HPS1 mice and at the 24-week timepoint for HPS2 mice. To account for the possibility of immune cells and other cell types present in HPS mice that could alter the total cell count and contribute to lower percentages of proSP-C^+^ cells, we quantified the percentage of proSP-C^+^ cells per total epithelial cells as denoted by NKX2.1 staining (**Supplemental Figure 2A-D**). There was a significant decrease in the percentage of proSP-C^+^, NKX2.1^+^ cells per total epithelial cells by NKX2.1 staining in HPS1 compared to WT mice at eight weeks of age (72.2 ± 3.2% vs. 78.2 ± 3.5, *p* = .046) and in HPS2 compared to WT mice at 24 weeks of age (65.0 ± 2.4% vs. 76.1 ± 3.9%, *p* = .003) (**Supplemental Figure 2A-D**). In contrast, in HPS3 mice, a subtype that has not been associated with pulmonary fibrosis, AT2 cell numbers were comparable to WT at eight and 24 weeks of age (**Supplemental Figure 3A, B**). Furthermore, in transgenic epithelial-corrected HPS2 (HPS2 TG) mice, AT2 cell numbers were significantly higher compared to HPS2 mice at 24 weeks of age (**Supplemental Figure 3A, C**).

Together, these data show that starting at eight weeks of age, prior to the onset of fibrosis, there is progressive loss of AT2 cells in HPS1, HPS2, and HPS1/2 mice, the HPS subtypes associated with fibrotic susceptibility (**Figure 1E**). We hypothesized that this loss could result from increased apoptosis, impaired proliferation, or aberrant differentiation (**Figure 1E**). Notably, no TUNEL-positive apoptotic AT2 cells were detected in WT or HPS mice across the same timepoints (**Supplemental Figure 4A**), suggesting that either AT2 cell apoptosis was a rare event in HPS mice, apoptotic cells were rapidly cleared, or loss of AT2 cells was occurring through predominantly through other mechanisms.

### Impaired AT2 cell response after acute influenza injury in HPS mice

To evaluate whether impaired proliferation contributes to AT2 cell loss, we employed an influenza model to evaluate AT2 cell function in the context of regeneration after physiological injury (**Figure 2A**) (14, 15). Because AT2 cell loss occurred in HPS1, HPS2, and HPS1/2 mice, we utilized HPS1 and HPS2 mice at eight weeks of age for the subsequent proliferation studies, as these mice best reflect human disease genetics. Weight change and oxygen saturation decreased similarly in WT, HPS1, and HPS2 mice during the first 10 days post-infection (dpi) with H1N1 PR8 influenza; subsequently, oxygen saturations trended lower in HPS1 and HPS2 mice though no mortality was observed in any group (**Figure 2B, C**). To ensure appropriate quantification and comparison across groups with the heterogeneous nature of influenza-induced lung injury, a computational image analysis approach was used to cluster image pixels based on H&E staining to define three regions: severely injured, mildly injured, and unaffected (**Supplemental Figure 5A**). There was no significant difference in the percentages of these regions or in the area formed by keratin 5 (KRT5)-expressing cells across all groups at 14 dpi (**Supplemental Figure 5A-E**).

**Figure 2.**
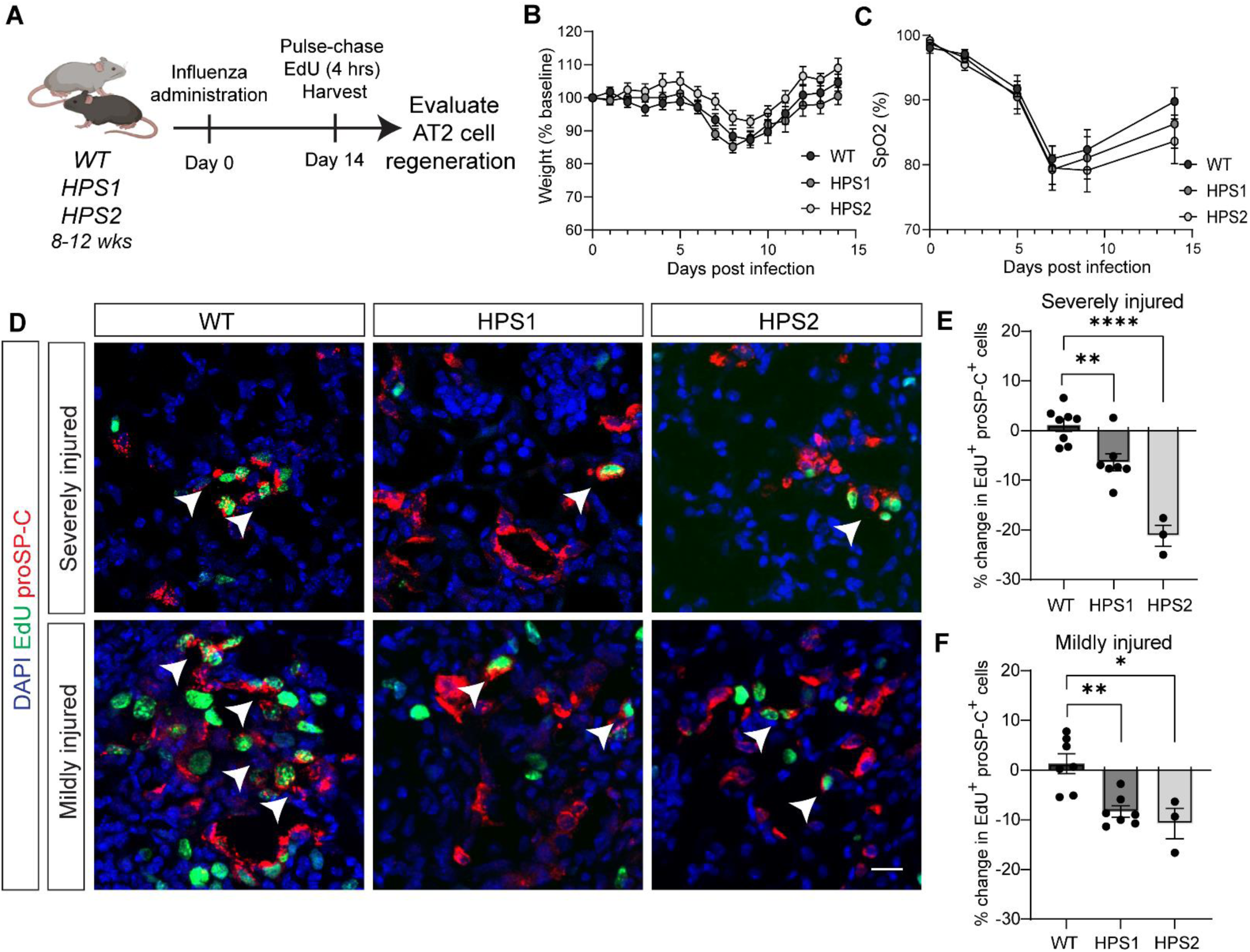
Impaired regeneration of HPS alveolar type II epithelial (AT2) cells after acute influenza injury. (A) Schematic of influenza experiment. (B, C) Mean change in (B) weight and (C) oxygen saturation (SpO_2_) in WT, HPS1, and HPS2 mice from 0 to 14 days post infection (dpi). (D) Immunofluorescence staining of lung tissue for proSP-C and EdU in WT, HPS1, and HPS2 mice at 14 dpi in severely and mildly injured regions. Arrows indicate proliferating AT2 cells (EdU^+^, proSP-C^+^). (E, F) Quantification of the percentage difference in proliferating AT2 cells (EdU^+^, proSP-C^+^) in (E) severely injured and (F) mildly injured regions at 14 dpi in HPS1 and HPS2 mice relative to WT. DAPI stains nuclei (blue). All quantification data are represented as mean ± SEM. Statistics using two-tailed unpaired Student’s *t* tests: adjusted * *p* < .05, ** *p* < .01, **** *p* < .0001 after Benjamini Hochberg correction for multiple comparisons. n = 3-10 per group. All scale bars, 20 μm. Schematic created with Biorender.com.

To examine epithelial regeneration after injury, we examined proliferating AT2 (EdU^+^, proSP-C^+^) cells in HPS1 and HPS2 as compared to WT mice after acute influenza infection (**Figure 2D-F**). The absolute numbers of EdU^+^, proSP-C^+^ cells were greatest in severely and mildly injured regions in WT mice; there were fewer EdU^+^, proSP-C^+^ cells in the corresponding regions in HPS1 and HPS2 mice (97.5 ± 21.9 EdU^+^, proSP-C^+^ cells in WT mice, 69.7 ± 30.8 in HPS1 mice, and 22.0 ± 5.6 cells in HPS2 mice for severely injured regions; 72.0 ± 5.1 EdU^+^, proSP-C^+^ cells in WT mice, 55.3 ± 23.0 in HPS1 mice, and 32.5 ± 1.0 cells in HPS2 mice in mildly injured regions). In unaffected areas where histologic evidence of injury was not observed, EdU^+^, proSP-C^+^ cells were extremely rare (not shown). Given the heterogeneous nature of influenza injury and previously observed AT2 cell loss in naïve HPS mice, we quantified the difference in the percentages of EdU^+^, proSP-C^+^ cells in WT, HPS1, and HPS2 mice. A decrease in the percentage of proliferating AT2 cells were observed in both severely injured and mildly injured regions in both HPS models at 14 dpi compared to WT (**Figure 2D, E, F**). In severely injured regions, there was a 6.4 ± 4.5% decrease in EdU^+^, proSP-C^+^ cells in HPS1 mice (adjusted *p* = .0033) and a 21.2 ± 3.7% decrease in HPS2 mice (adjusted *p* < .0001) compared to WT (**Figure 2D, E**). In mildly injured regions, there was an 8.3 ± 3.0% reduction in EdU^+^, proSP-C^+^ cells in HPS1 mice (adjusted *p* = .0012) and a 10.8 ± 5.3% decrease in HPS2 mice (adjusted *p* = .011) compared to WT (**Figure 2D, F**). Overall, these data demonstrate impaired HPS AT2 cell proliferation in the setting of acute influenza injury.

### Intrinsic HPS AT2 cell proliferation defect ex vivo and in vivo

To evaluate whether this impaired HPS AT2 cell proliferation was due to cell-intrinsic defects in progenitor function and proliferative capacity, we first employed an *ex vivo* organoid model (15). We generated lung organoids with WT primary outgrowth lung fibroblasts and AT2 cells from WT, HPS1, and HPS2 mice at eight weeks of age (**Figure 3A**). Compared to lung organoids generated with WT AT2 cells, those with HPS2 AT2 cells were significantly smaller in size (adjusted *p* = .002) and displayed significantly decreased colony-forming efficiency (CFE), 2.5 ± 0.4% compared to 4.2 ± 0.4% in WT organoids (adjusted *p* = .043), over 21 days of culture (**Figure 3B-E**). In contrast, organoids generated from HPS1 AT2 cells demonstrated comparable size and growth compared to those generated with WT AT2 cells (**Figure 3B-E**).

**Figure 3.**
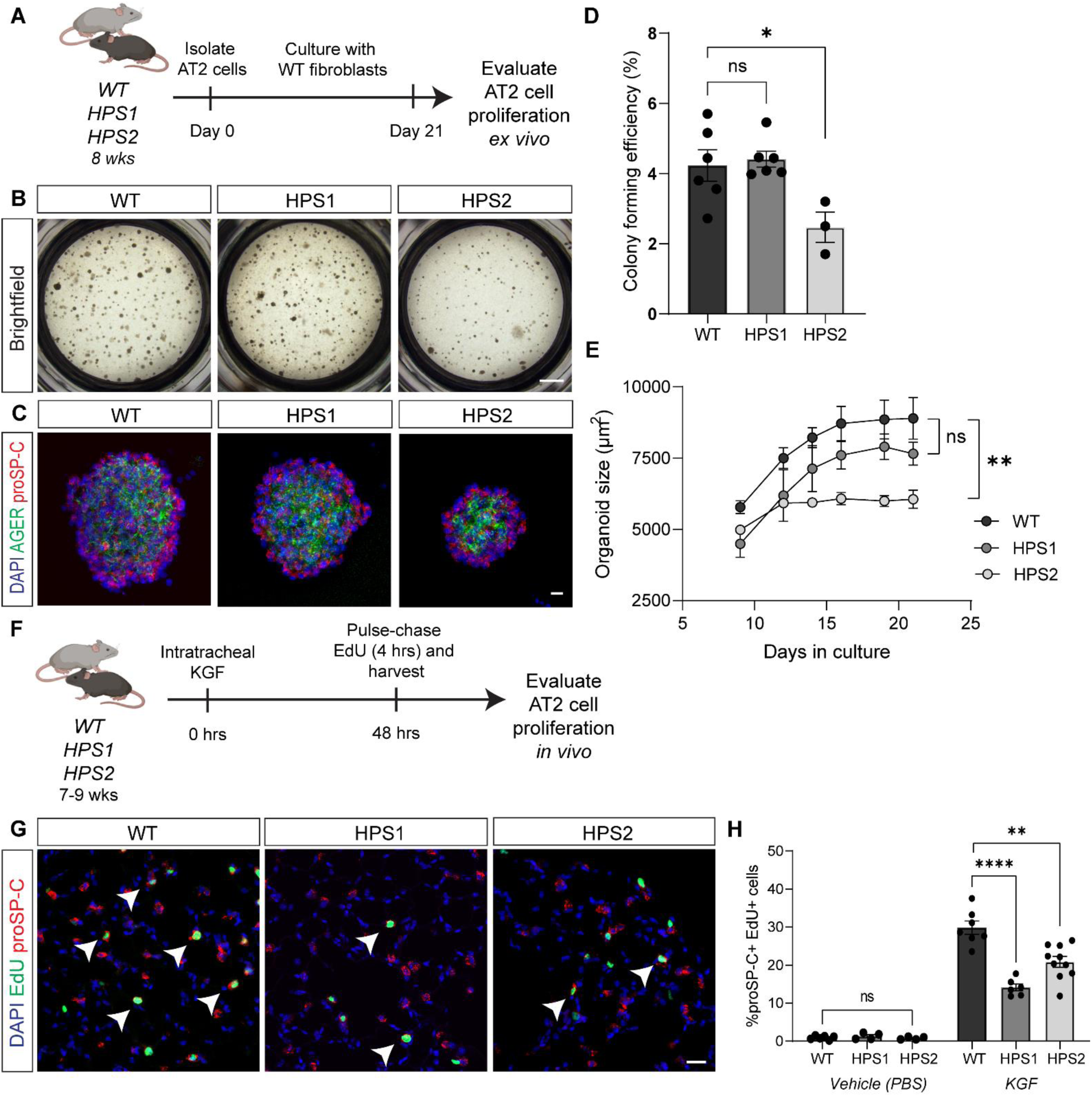
Impaired proliferation of HPS alveolar type II epithelial (AT2) cells *ex vivo* and *in vivo*. (A) Schematic of lung organoid culture system. (B) Brightfield imaging and (C) immunofluorescence (IF) staining of lung organoids for AGER and proSP-C generated with AT2 cells isolated from WT, HPS1, or HPS2 mice with WT fibroblasts at day 21 of culture. (D) Quantification of WT, HPS1, and HPS2 organoid colony forming efficiency (CFE) at day 21 of culture. (E) Quantification of WT, HPS1, and HPS2 organoid size from day 9 to day 21 of culture. (F) Schematic of keratinocyte growth factor (KGF) experiment. (G) IF staining of lung tissue for proSP-C and EdU in WT, HPS1, and HPS2 mice 48 hours after KGF administration. Arrows indicate proliferating AT2 cells (EdU^+^, proSP-C^+^). (H) Quantification of the percentage of proliferating AT2 cells (EdU^+^, proSP-C^+^) in WT, HPS1, and HPS2 mice treated with vehicle (PBS) vs. KGF. DAPI stains nuclei (blue). All quantification data are represented as mean ± SEM. Statistics using two-tailed unpaired Student’s *t* tests: ns, not significant; adjusted * *p* < .05, ** *p* < .01, **** *p* < .0001 after Benjamini Hochberg correction for multiple comparisons. For (D-E), each point represents the average of at least three replicate wells from one mouse for a total of n = 3-6 mice per group. For (H), n = 4-10 per group per treatment. Scale bars in (B) 1 mm; in (C, G) 20 μm. Schematics created with Biorender.com.

To evaluate intrinsic AT2 cell proliferative capacity *in vivo*, we administered an epithelial cell-specific mitogen, keratinocyte growth factor (KGF), to WT, HPS1, and HPS2 mice at eight weeks of age and quantified the percentage of EdU^+^, proSP-C^+^ cells 48 hours after KGF administration (**Figure 3F**). The percentage of proliferating AT2 cells was significantly reduced in HPS1 (14.2% EdU^+^, proSP-C^+^ cells, adjusted *p* < .0001) and HPS2 mice (20.9 ± 4.5% EdU^+^, proSP-C^+^ cells, adjusted *p* = 0.001) compared to WT (29.9 ± 4.6% EdU^+^, proSP-C^+^ cells) (**Figure 3G, H**). Low rates of AT2 cell proliferation were observed in WT, HPS1, and HPS2 mice administered vehicle (**Figure 3H**). Overall, these data demonstrate an intrinsic proliferative defect in HPS AT2 cells *ex vivo* and *in vivo*.

### Lineage tracing of HPS AT2 cells reveals spontaneous aberrant differentiation at 8 weeks of age

To confirm the finding of loss of AT2 cells identified based on proSP-C expression and further investigate the etiology of AT2 cell loss by evaluating cell fate and differentiation, we utilized the *Sftpc*^CreERT2/+^;*R26R*^EYFP/+^ reporter to generate four additional lines of mice (WT, HPS1, HPS2, and HPS1/2 mice, each with *Sftpc*^CreERT2/+^;*R26R*^EYFP/+^) to perform lineage tracing of AT2 cells. Recombination with tamoxifen was induced at four weeks of age, prior to the timepoint where we observed onset of significant AT2 cell loss based on proSP-C immunostaining (**Figure 4A**). Quantification of the number of lineage-labeled AT2 cells (EYFP^+^) in paraffin-embedded sections confirmed significant loss of AT2 cells starting at eight weeks of age in HPS1, HPS2, and HPS1/2 mice, with quantitative results similar to those obtained using proSP-C immunostaining (**Supplemental Figure 6A, B**). HPS1/2 *Sftpc*^CreERT2/+^;*R26R*^EYFP/+^ mice exhibited 9.0 ± 0.8% lineage-labeled AT2 cells compared to 14.4 ± 0.8% in WT *Sftpc*^CreERT2/+^;*R26R*^EYFP/+^ mice (adjusted *p* < 0.001) at eight weeks of age (**Supplemental Figure 6A, B**).

**Figure 4.**
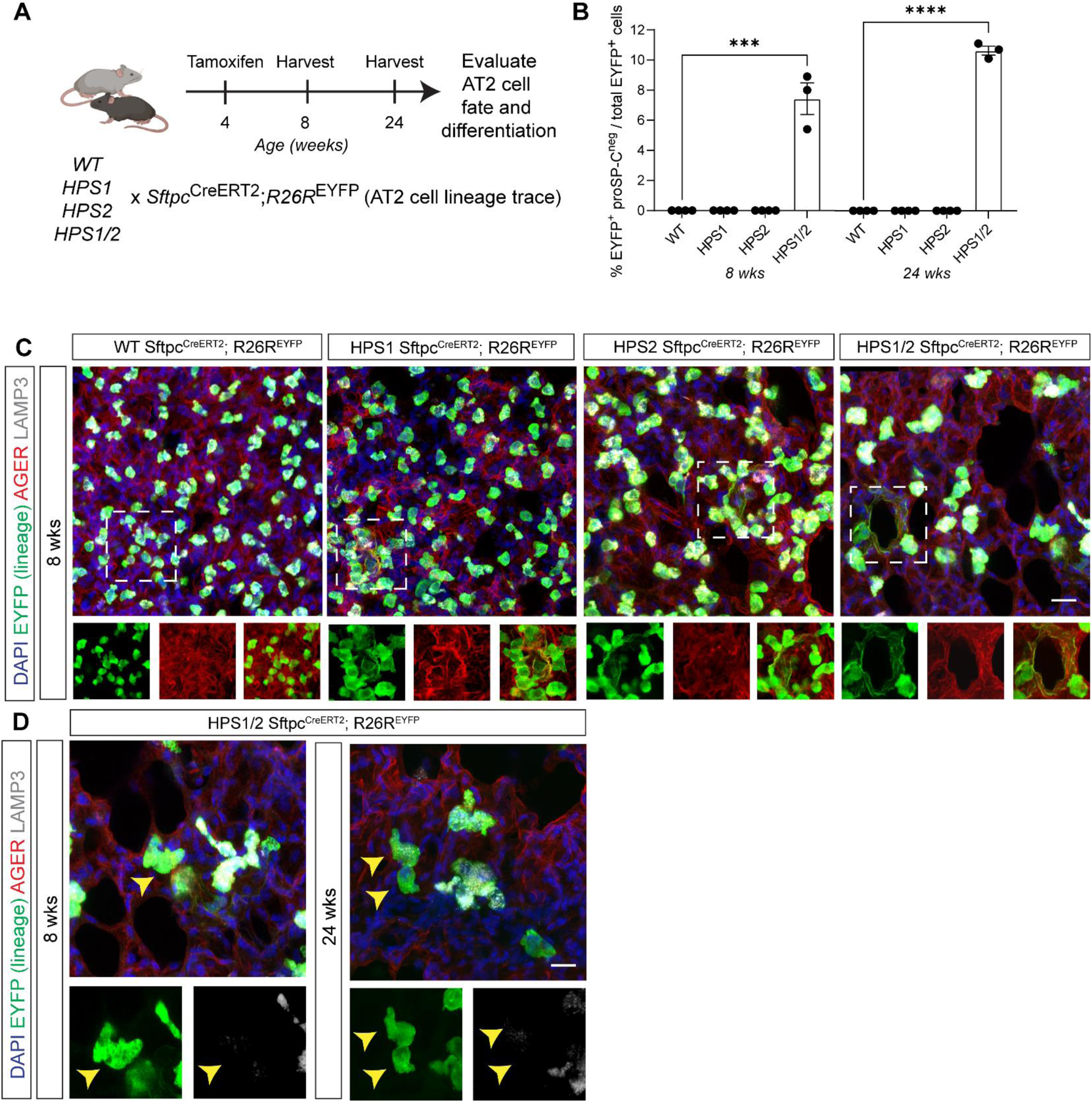
Lineage tracing of alveolar type II epithelial (AT2) cells in HPS mice demonstrates aberrant differentiation at 8 weeks of age. A) Schematic of lineage labeling AT2 cells in WT, HPS1, HPS2, and HPS1/2 *Sftpc*^CreERT2/+^;*R26R*^EYFP/+^ mice. (B) From images in Supplemental Figure 6A, quantification of EYFP^+^ proSP-C^neg^ cells as a percentage of EYFP^+^ lineage cells in WT, HPS1, HPS2, and HPS1/2 *Sftpc*^CreERT2/+^;*R26R*^EYFP/+^ mice at 8 and 24 weeks of age. (C) Immunofluorescence (IF) staining of whole-mount lung tissue for EYFP, AGER, and LAMP3 in WT, HPS1, HPS2, and HPS1/2 *Sftpc*^CreERT2/+^;*R26R*^EYFP/+^ mice at 8 weeks of age. Dashed line boxes represent squamous EYFP^+^, AGER^+^, LAMP3^neg^ AT2-derived alveolar type I (AT1) cells with AGER and EYFP and merged images shown below. (D) IF staining of HPS1/2 *Sftpc*^CreERT2/+^;*R26R*^EYFP/+^ mice at 8 and 24 weeks of age, with arrows indicating cuboidal EYFP^+^, AGER^neg^, LAMP3^neg^ cells with EYFP and LAMP3 images shown below. HPS1/2 *Sftpc*^CreERT2/+^;*R26R*^EYFP/+^ 8 week images in (C) and redemonstrated again in (D) capture simultaneous presence of squamous EYFP^+^, AGER^+^, LAMP3^neg^ and cuboidal EYFP^+^, AGER^neg^, LAMP3^neg^ cells. DAPI stains nuclei (blue). All quantification data are represented as mean ± SEM. Statistics using two-tailed unpaired Student’s *t* tests: *** *p* < .001; **** *p* < .0001. n = 3-4 per group per time point. Scale bars in (C, D), 20 μm (inset boxes equal scale to main image). Schematic created with Biorender.com.

Loss of lineage-labeled AT2 cells was also seen at eight weeks of age in HPS1 *Sftpc*^CreERT2/+^;*R26R*^EYFP/+^ mice (11.8 ± 0.2% EYFP^+^ cells, adjusted *p* = .003) and HPS2 *Sftpc*^CreERT2/+^;*R26R*^EYFP/+^ (12.9 ± 1.1% EYFP^+^ cells, adjusted *p* = .034) compared to WT *Sftpc*^CreERT2/+^;*R26R*^EYFP/+^ mice (**Supplemental Figure 6A, B**).

In HPS1/2 *Sftpc*^CreERT2/+^;*R26R*^EYFP/+^ mice, we observed the emergence of lineage-labeled cells that did not express proSP-C (EYFP^+^, proSP-C^neg^), constituting 7.4 ± 1.8% of all lineage-labeled cells at 8 weeks of age and 10.6 ± 0.5% at 24 weeks of age (**Figure 4B; Supplemental Figure 6A**). In contrast, EYFP^+^, proSP-C^neg^ cells were not identified in WT, HPS1, or HPS2 *Sftpc*^CreERT2/+^;*R26R*^EYFP/+^ mice at 8 or 24 weeks of age (**Figure 4B; Supplemental Figure 6A**). We next proceeded to investigate the identity of these lineage-labeled cells and AT2-to-AT1 cell differentiation as a potential etiology of AT2 cell loss. To account for the squamous morphology of AT1 cells, we performed immunostaining of thicker 150 μm whole-mount lung sections. Utilizing this technique, lineage-labeled EYFP^+^, AGER^+^, LAMP3^neg^ cells with the squamous morphology of AT1 cells were identified in the lungs of HPS1, HPS2, and HPS1/2 *Sftpc*^CreERT2/+^;*R26R*^EYFP/+^ mice, starting at eight weeks of age (**Figure 4C**). No lineage-labeled AT1 cells were detected in WT *Sftpc*^CreERT2/+^;*R26R*^EYFP/+^ mice (**Figure 4C**). Notably, cuboidal lineage-labeled EYFP^+^, AGER^neg^, LAMP3^neg^ cells were also observed, but only in HPS1/2 *Sftpc*^CreERT2/+^;*R26R*^EYFP/+^ mice, starting at eight weeks of age and persisting through 24 weeks of age (**Figure 4D**). These lineage tracing results confirm AT2 cell loss and reveal spontaneous and aberrant differentiation of AT2 cells in HPS mice starting at eight weeks of age.

### Transcriptomic signatures of senescence and aberrant differentiation are enriched in HPS AT2 cells

Given the aberrant differentiation and impaired proliferation exhibited by HPS AT2 cells, we sought to identify pathways that could underlie impaired AT2 cell progenitor function and connect them with mechanisms known to drive pulmonary fibrosis. As we observed these changes in progenitor cell function early in the disease course, prior to the onset of fibrosis, we examined the transcriptomes of AT2 cells isolated from WT, HPS1, HPS2, and HPS1/2 mice at eight weeks of age (**Figure 5A**). Fluorescence activated cell sorting (FACS) of AT2 cells revealed a significant decrease in the percentage of AT2 cells captured from HPS mice compared to WT (**Supplemental Figure 7A-D**). Principal component analysis (PCA) revealed significantly different transcriptomes in AT2 cells from WT, HPS1, HPS2, and HPS1/2 mice (**Figure 5B**). Differential gene expression testing and over-representation analysis (ORA) demonstrated upregulation of genes associated with immune cell activation and cell cycle regulation in HPS AT2 cells (**Supplemental Figure 8A-J**). HPS AT2 cells displayed decreased expression of canonical markers of AT2 cells, increased expression of AT1 cell markers, as well an increase in expression of genes associated with a reprogramed transitional cell state, including *Krt8*, *Cldn4*, *Lgals3, and Krt19* (**Figure 5C**) (16–18). These observations prompted us to examine markers of p53 signaling and cellular senescence, including *Trp53*, *Cdkn1a*, *Cdkn2d*, and *Ccnd1*, which were upregulated in HPS1, HPS2, and HPS1/2 AT2 cells, with the greatest change in expression seen in HPS1/2 AT2 cells (**Figure 5C, D**) (19–21). In addition, by 48 weeks of age, when significant fibrotic disease is present, 5.9 ± 1.8% of AT2 cells isolated from HPS1/2 mice stained positive for β-galactosidase, a marker for senescent cells. In contrast, β-galactosidase activity was not detected in AT2 cells from WT or HPS mice at 8 weeks of age or WT, HPS1, or HPS2 mice at 48 weeks of age (*p* = .005) (**Figure 5E, F**).

**Figure 5.**
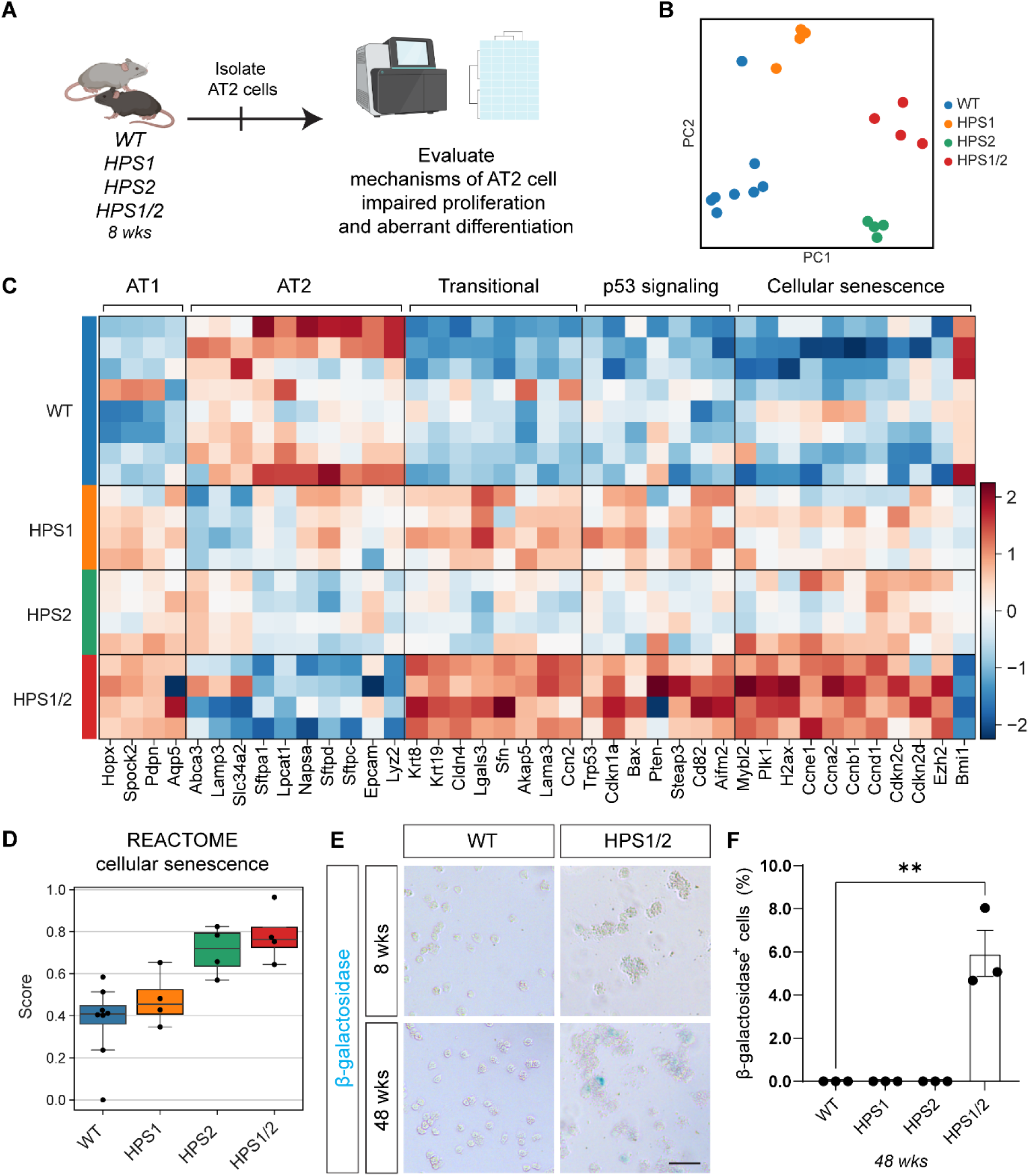
Transcriptomic analysis suggests aberrant differentiation and p53-mediated senescence in HPS alveolar type II epithelial (AT2) cells. (A) Experimental schematic for bulk mRNA sequencing of HPS AT2 cells. (B) Principal component analysis (PCA) of AT2 cells from WT, HPS1, HPS2, and HPS1/2 mice at 8 weeks of age. (C) Differential expression analysis of AT2, transitional, and AT1 cell genes and p53 signaling pathway and cellular senescence genes in AT2 cells from WT, HPS1, HPS2, and HPS1/2 mice at 8 weeks of age. (D) Overall gene set score for cellular senescence. (E) Representative β-galactosidase staining of AT2 cells isolated from WT and HPS1/2 mice at 8 and 48 weeks of age. (F) Quantification of the percentage of β-galactosidase^+^ cells in WT, HPS1, HPS2, and HPS1/2 mice at 48 weeks of age. DAPI stains nuclei (blue). All quantification data are represented as mean ± SEM. Statistics using two-tailed unpaired Student’s *t* tests: ** *p* < .01. n = 4-8 per group for bulk mRNA sequencing, n = 3 per group for β-galactosidase staining. Scale bars in (E), 50 μm.

### HPS AT2 cells are primed to enter a p53-mediated Krt8^+^ reprogrammed transitional cell state, which are present in lung tissue from a patient with HPS-1

With the aberrant differentiation exhibited by HPS AT2 cells and transcriptomic evidence of a reprogrammed transitional cell state, we sought to identify this cell state *in situ*. In HPS1/2 *Sftpc*^CreERT2/+^;*R26R*^EYFP/+^ mice, the previously captured lineage-labeled cuboidal EYFP^+^, LAMP3^neg^ cells (**Figure 4D**) demonstrated co-expression of markers of a reprogrammed transitional cell state, including KRT8, CLDN4, LGALS3, and KRT19 starting at eight weeks and accumulating through 24 weeks of age (**Figure 6A, B**; **Supplemental Figure 9A-C**). While KRT8^+^ cells were not detected in WT, HPS1, or HPS2 *Sftpc*^CreERT2/+^;*R26R*^EYFP/+^ mice, KRT19^+^ cells were present in HPS1 and HPS2 *Sftpc*^CreERT2/+^;*R26R*^EYFP/+^ mice at eight weeks of age (**Figure 6A, B; Supplemental Figure 9D**). In all HPS models, squamous lineage-labeled EYFP^+^, LAMP3^neg^ cells did not express KRT19, suggesting they were AT2-derived AT1 cells rather than reprogrammed transitional cells (**Figure 6B**).

**Figure 6.**
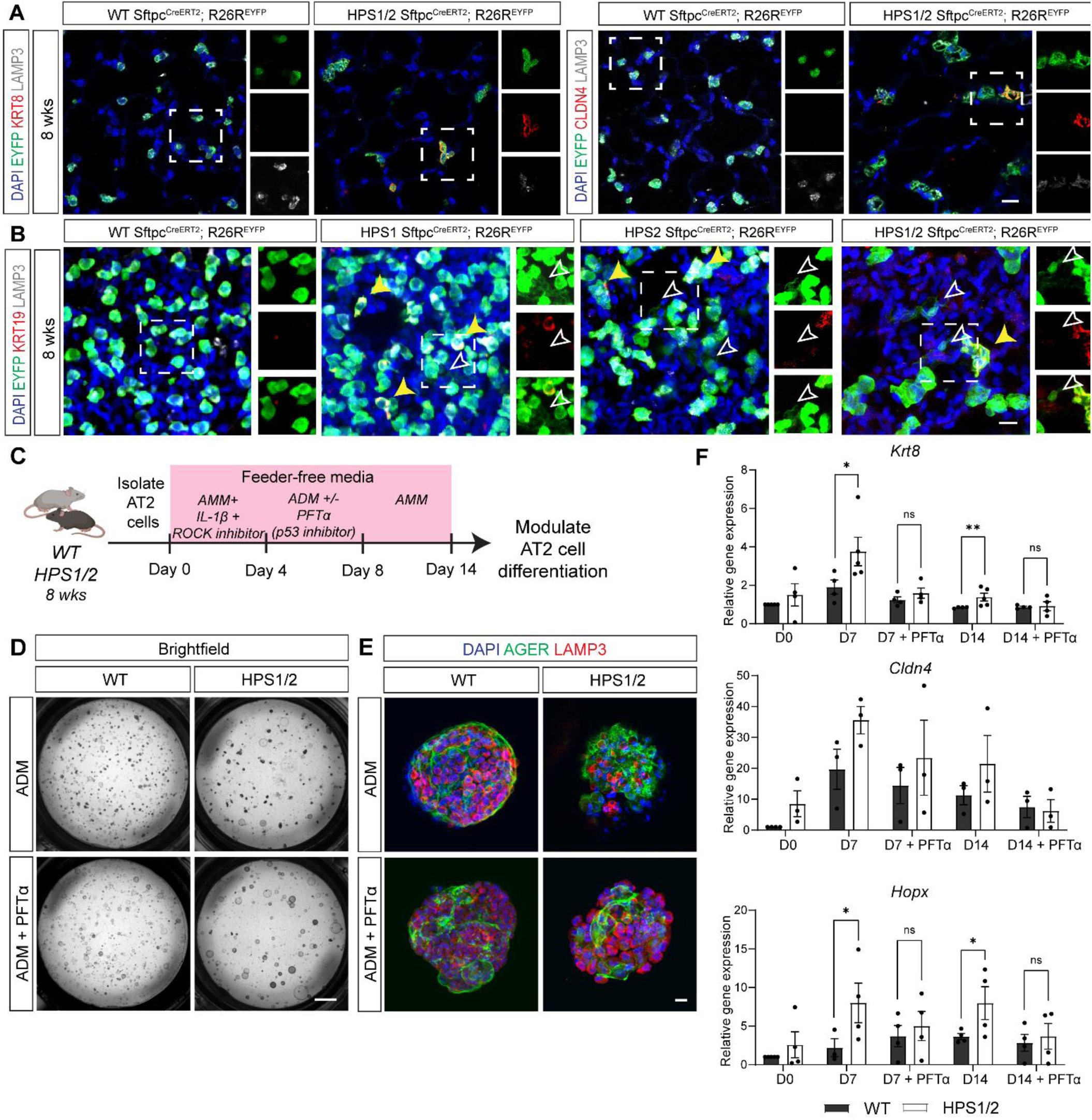
A p53-mediated Krt8^+^ reprogrammed transitional cell state in HPS mice. (A) Immunofluorescence (IF) staining of paraffin-embedded lung tissue for EYFP, KRT8 or CLDN4, and LAMP3 in WT and HPS1/2 *Sftpc*^CreERT2/+^;*R26R*^EYFP/+^ mice at 8 weeks of age. Dashed line boxes represent EYFP^+^ cells with inset boxes displaying EYFP, KRT8 or CLDN4, and LAMP3 staining. (B) IF staining of whole-mount lung tissue for EYFP, KRT19, and LAMP3 in WT, HPS1, HPS2, and HPS1/2 *Sftpc*^CreERT2/+^;*R26R*^EYFP/+^ mice at 8 weeks of age. White arrows indicate squamous EYFP^+^, KRT19^neg^, LAMP3^neg^ AT2-derived alveolar type I (AT1) cells and yellow solid arrows indicate cuboidal EYFP^+^, KRT19^+^ cells; dashed line boxes and insets display EYFP and KRT19 and merged images. (C) Schematic for alveolosphere culture system with feeder-free alveolar maintenance media (AMM) and alveolar differentiation media (ADM) and treatment with pifithrin-α (PFTα). (D) Brightfield imaging and (E) IF staining of alveolospheres for AGER and LAMP3 generated with alveolar epithelial type II (AT2) cells from WT or HPS1/2 mice with and without treatment with PFTα at day 14 (D14) of culture. (F) Relative gene expression of selected reprogrammed transitional cell (*Krt8*, *Cldn4*) and alveolar epithelial type I (AT1) cell (*Hopx*) genes in alveolospheres at day 0 (D0), day 7 (D7), and day 14 (D14) of culture with and without treatment with PFTα. Relative gene expression compared to WT at D0. Each point represents four replicate wells per time point for a total of n = 3-4 mice. DAPI stains nuclei (blue). All quantification data are represented as mean ± SEM. Statistics using two-tailed unpaired Student’s *t* tests: ns, not significant; * *p* < .05; ** *p* < .01. Scale bars in (A, B, E), 20 μm (inset boxes equal scale to main image); (D) 1 mm. Schematic created with Biorender.com.

To further investigate the dynamics of AT2 cell differentiation and of this Krt8^+^ reprogrammed transitional cell state, we utilized an *in vitro* feeder-free alveolosphere culture system to evaluate cell intrinsic differentiation (**Figure 6C**) (22, 23). Alveolospheres were generated with AT2 cells from WT, HPS1, HPS2, and HPS1/2 mice at eight weeks of age, producing AT1 cells as indicated by AGER staining (**Figure 6D, E**). Upon induction of differentiation, *Krt8*, *Cldn4*, and *Hopx* expression increased across all groups. Expression of *Krt8* and *Hopx* was significantly higher in HPS1/2 alveolospheres compared to WT on day 7 and continued to be significantly upregulated compared to WT on day 14 (**Figure 6F**). As transcriptomic data suggested this aberrant differentiation could be driven by p53 activation, alveolospheres were treated with pifithrin-α (PFTα), a p53 inhibitor, during induction of differentiation. Treatment with PFTα decreased *Krt8* and *Hopx* expression in HPS1/2 alveolospheres on day 7 and day 14 of culture to levels comparable to those in WT alveolospheres (**Figure 6F**). Similar trends in gene expression were seen in HPS1 and HPS2 alveolospheres compared to WT (**Supplemental Figure 10B-D**). Expression of *Sftpc* was suppressed across all groups in alveolosphere culture; with induction of differentiation, increases in *Sftpc* expression after treatment with PFTα were not statistically significant (**Supplemental Figure 10A**).

Finally, explanted lung tissue from a patient with HPS-1 pulmonary fibrosis revealed proSP-C^+^ AT2 cells that stained positive for β-galactosidase (**Figure 7A**). KRT8-expressing cells which co-expressed an epithelial cell marker, NKX2.1, were also detected (**Figure 7B**). A subset of these KRT8^+^, NKX2.1^+^ cells also expressed HTII-280, a human AT2 cell marker (**Figure 7B**). Neither β-galactosidase^+^ AT2 cells nor KRT8^+^ epithelial cells were identified in lung tissue from age-matched and sex-matched control donor patients (**Figure 7A, B**). Taken together, these results demonstrate that HPS AT2 cells are primed to differentiate and persist in a Krt8^+^ reprogrammed transitional cell state, which can be ameliorated *in vitro* in HPS1/2 mice by inhibiting the activity of p53.

**Figure 7.**
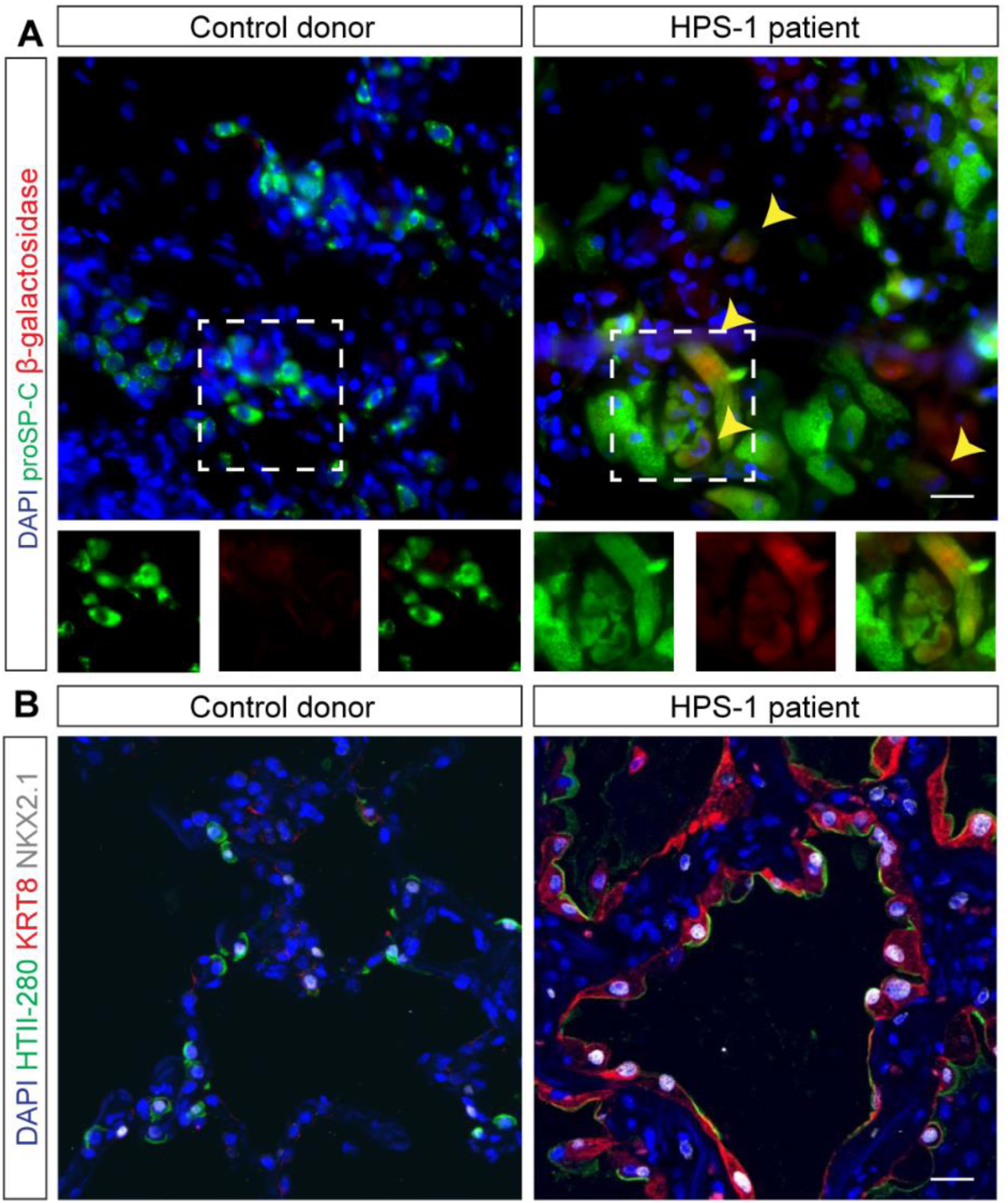
Senescence and a Krt8^+^ reprogrammed transitional cell state in a patient with HPS-1. (A) Immunofluorescence (IF) staining of paraffin-embedded lung tissue from an age- and sex-matched control donor patient and HPS-1 patient for proSP-C and β-galactosidase. Arrows and dashed line boxes represent proSP-C^+^ cells with inset boxes displaying proSP-C and β-galactosidase staining with merged images. (B) IF staining of paraffin-embedded lung tissue from an age- and sex-matched control donor patient and HPS-1 patient for HTII-280, KRT8, and NKX2.1. DAPI stains nuclei (blue). All scale bars, 20 μm (inset boxes equal scale to main image).

## Discussion

As a monogenic disorder with murine models that recapitulate key features of the disease, HPS presents a unique opportunity for the study of progressive pulmonary fibrosis. Although the lysosomal and endosomal protein trafficking defects in HPS have been well-characterized, the link between these dysregulated cellular processes and the downstream functional consequences for the alveolar epithelium and their role in fibrotic remodeling remains incompletely understood. In addition, there has been debate about whether mechanisms in HPS-PF are distinct from other heritable forms of pulmonary fibrosis and IPF (24, 25). In this study, we demonstrate early disruption of alveolar epithelial maintenance in HPS in the setting of AT2 cell progenitor dysfunction. We propose that over time, impaired AT2 cell proliferation and aberrant differentiation contribute to progressive AT2 cell loss, and ultimately, senescence and depletion of the stem cell pool with pulmonary fibrosis (**Figure 8**). Intrinsic AT2 progenitor cell dysfunction is a novel mechanism in HPS-PF primed for early intervention and highlights convergent mechanisms of disease between HPS-PF and other forms of pulmonary fibrosis.

**Figure 8.**
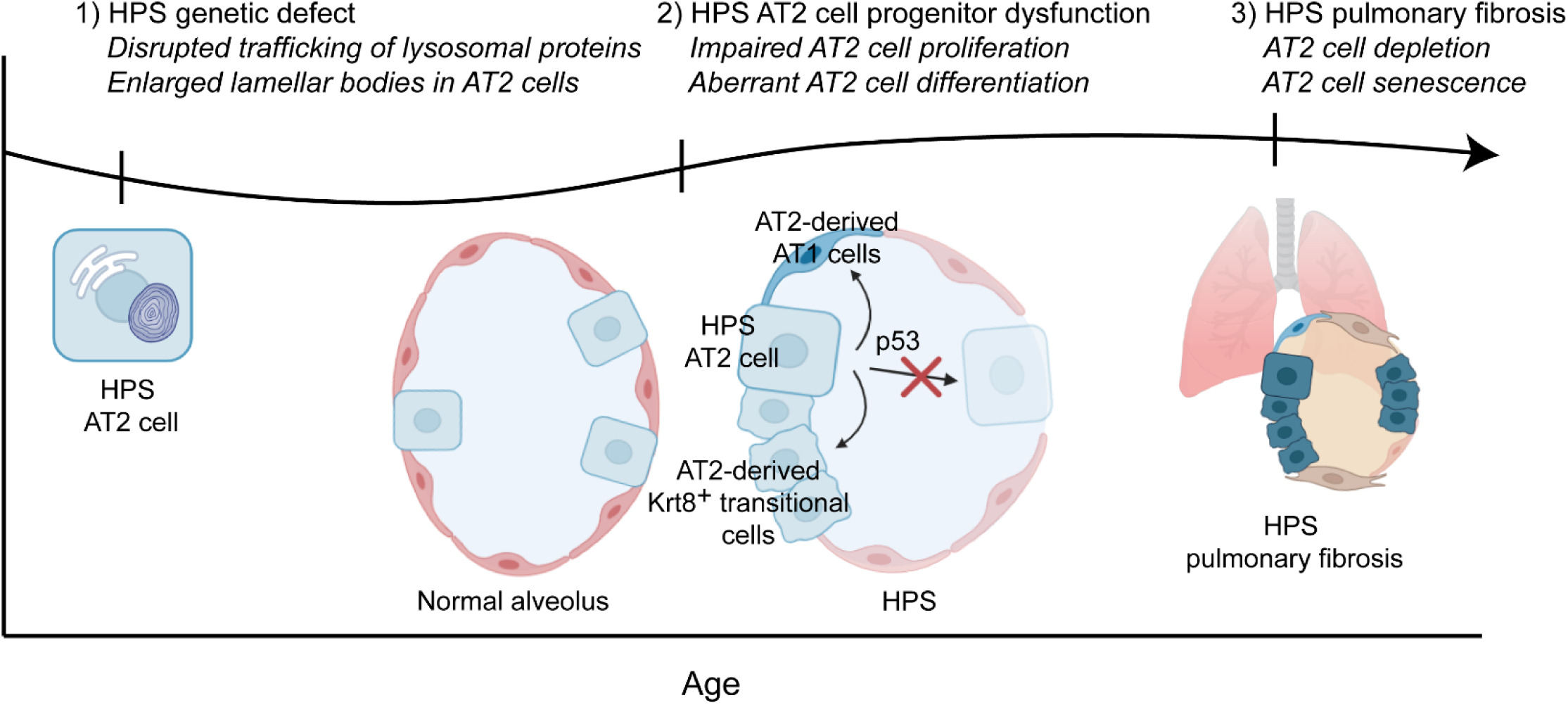
Time course of alveolar type II epithelial (AT2) cell dysfunction in HPS pulmonary fibrosis. Our current working theory is that the HPS genetic mutations disrupt AT2 progenitor cell function early, with activation of the p53 pathway, impairing AT2 cell proliferation and driving aberrant AT2 cell differentiation. Over time, this results in accelerated AT2 cell loss with senescence and depletion of the stem cell pool and progressive fibrotic remodeling.

Under homeostatic conditions, AT2 cells are largely quiescent yet some degree of proliferation is required to maintain the alveolar epithelium (26). Other models have demonstrated that interruption of AT2 cell proliferative capacity contributes to fibrotic remodeling (19, 27). While we observed similarly low rates of AT2 proliferation in WT and HPS mice under basal conditions, the decreased rates of proliferation in HPS AT2 cells were striking in multiple experimental conditions including in an *ex vivo* lung organoid model and in two *in vivo* models, after mitogenic stimulation and with an influenza injury model. This impaired proliferation may result from not only depletion of the AT2 cell progenitor pool, but also decreased proliferative capacity of these progenitor cells as a result of the HPS mutations.

Notably, HPS1 AT2 cells appeared to perform comparably to WT AT2 cells in an organoid model but exhibited impaired proliferation after administration of KGF and after injury in an influenza model. This may reflect differences in the consequences of the HPS1 and HPS2 mutations on AT2 cell function as well as the models utilized to study cell proliferation and regeneration (10, 28). Furthermore, differences in AT2 cell regenerative capacity by injury severity after acute influenza infection suggests that the HPS mutations have divergent effects on AT2 cell subpopulations, particularly those that function as progenitor cells (15, 29).

In addition to impaired proliferation, HPS AT2 cell progenitor dysfunction appears to also manifest with aberrant differentiation. In adults, AT2-to-AT1 cell differentiation is typically observed in the setting of injury, as a subset of progenitor cells are mobilized to regenerate the alveolar epithelium (15, 30). This process occurs at very low rates during homeostasis, with an approximate turnover rate of 0.005% of the alveolar surface with new AT2 cell-derived AT1 cells per day (31). In bleomycin models of pulmonary fibrosis, epithelial cell subpopulations enriched in expression of *Krt8*, *Cldn4*, *Lgals3*, and *Krt19*, which have been referred to as pre-alveolar type 1 transitional cell state (PATS), damage-association transition progenitors (DATPs), and Krt8^+^ alveolar differentiation intermediates (ADIs), appear to emerge from incomplete differentiation of AT2 cells (16–18). In HPS, through lineage tracing and alveolosphere studies without exogenous stimuli, we demonstrate that AT2 cells from HPS1, HPS2, and HPS1/2 mice display aberrant differentiation. The most severe phenotype with the emergence of a Krt8^+^ reprogrammed transitional cell state was observed in HPS1/2 mice, a model system associated with spontaneous fibrosis, underscoring the potential role of these cells in fibrotic remodeling. Our data further supports emerging evidence that this state may not necessarily be “transitional” in the sense that cells may lose their capacity to differentiate into AT1 cells. For some cells, entry into this state may be unidirectional, resulting in loss of the ability to contribute to functional alveolar epithelial maintenance and repair (32).

Given the widespread and nonselective injury induced by bleomycin, the contribution of other cell types and potential AT1 cell dysfunction in driving aberrant AT2 cell differentiation and the emergence of the Krt8^+^ reprogrammed transitional cell state has been debated (33). In subsequent studies, this cell state has been identified in genetic models with selective AT2 cell modification, including models of accelerated senescence of AT2 cells via *Trp53* activation, surfactant metabolism dysfunction (*Sftpc*^C185G^mice), and Nkx2.1 ablation in alveolar epithelial progenitors (19, 34, 35). Here, in a feeder-free alveolosphere culture system with only AT2 cells and no stromal support cells, HPS AT2 cells appear primed to differentiate and persist in this Krt8^+^ reprogrammed transitional cell state. Taken together, this suggests that aberrant differentiation in HPS is driven by intrinsic AT2 cell dysfunction, although other cell types may provide additional feedback to further amplify this process (32).

In the setting of impaired progenitor function and progressive AT2 cell loss, transcriptomic analyses highlighted a role for p53 activation in driving AT2 cell dysfunction and pulmonary fibrosis in HPS. From the study of telomere disorders, p53-mediated senescence of AT2 cells has long been implicated in pulmonary fibrosis (36). In an experimental murine model, conditional loss of function of SIN3 transcription regulator homolog A (*Sin3a*), a regulator of Trp53 acetylation, in AT2 cells induced senescence, impaired proliferation, and caused loss of AT2 cells, resulting in spontaneous, progressive fibrosis. Mice with loss of function of both *Sin3a* and p53 did not exhibit AT2 cell loss and did not develop significant fibrosis, indicating that activation of the p53 pathway is necessary to induce AT2 cell senescence and fibrosis (19). Previous studies of the Krt8^+^ transitional reprogrammed cell state have demonstrated that these cells show increased expression of genes involved in p53, DNA damage, and cellular senescence pathways (17, 19, 32). Indeed, inhibition of p53 was sufficient to attenuate upregulation of *Krt8* and *Hopx* in alveolospheres generated with HPS AT2 cells compared to WT, suggesting that p53 activation is the driver of aberrant differentiation and this transitional cell state in HPS. Prior studies have demonstrated that HPS AT2 cells have excess production of monocyte MCP-1, a senescence-associated secretory phenotype (SASP) factor, and epithelial-specific deletion of MCP-1 reduced fibrotic susceptibility in HPS mice (12, 37). HPS-1 patients have also been shown to have elevated levels of MCP-1 in bronchoalveolar lavage (BAL) fluid that correlate with lung disease severity (38). Additional studies are required to evaluate the secretome of HPS AT2 cells, as well as further examine interactions between AT2 cells, macrophages, and fibroblasts, to identify which factors drive fibrotic remodeling in HPS.

Although increased apoptosis of AT2 cells has been observed across many forms of pulmonary fibrosis, there remains debate about whether depletion of AT2 cells is necessary or sufficient to drive fibrotic remodeling (25, 39–41). Using diphtheria toxin-mediated epithelial cell ablation in SPC-DTR (diphtheria toxin receptor) mice, Sisson et al. showed that AT2 cell loss resulted in fibrotic remodeling. When Yao et al selectively ablated AT2 cells in *Sftpc*^CreER^;*Rosa*^DTA^ ;*Rosa*^mTmG^ mice, significant AT2 cell loss and collagen deposition were observed, but fibrosis diminished over time, suggesting a transient effect. In contrast, progressive pulmonary fibrosis was observed in *Sin3a* loss of function mice with AT2 cell senescence in addition to AT2 cell loss (19). Although previous studies have described increased AT2 cell apoptosis in HPS1/2 mice as well as in bleomycin-challenged but not naïve HPS1 and HPS2 single mutant mice, we did not detect AT2 cell apoptosis in any of the HPS models up to 48 weeks of age (8, 10). In addition to differences in techniques and assays used to evaluate apoptosis, our ability to capture rare events may have been limited without a stereotactic approach, and we cannot rule out that other mechanisms of increased cell death, including necroptosis, may also contribute to AT2 cell loss in HPS.

The exact mechanisms by which the HPS endosomal trafficking mutations result in p53 activation and AT2 progenitor cell dysfunction are unknown. There have been several prior reports demonstrating the emergence of a reprogrammed transitional cell state with lung injury and with disrupted proteostasis (42–46). While the HPS mutations could produce cell cycle defects that directly result in cell cycle arrest, it is also possible that they contribute to other forms of cellular dysfunction, including mitochondrial dysfunction and aberrant autophagy, which can induce cellular senescence (6, 9). As these mechanisms have been implicated in other fibrotic lung diseases, there may be interconnecting pathways contributing to AT2 cell dysfunction in HPS. In mice, there may be an additive effect of the HPS1 and HPS2 mutations that results in markedly impaired progenitor cell function and accelerated senescence with the emergence of a Krt8^+^ reprogrammed transitional cell state in double mutant HPS1/2 AT2 cells. This could underlie the divergent phenotypes observed, with fibrotic sensitivity in HPS1 and HPS2 mice in comparison to the development of spontaneous, progressive fibrosis in HPS1/2 mice. It is important to note, however, that patients only bear single variants, either in *HPS1* or *HPS2.* The emergence of KRT8^+^ epithelial cells and spontaneous pulmonary fibrosis in human HPS patients is likely to result from the combined effects of genetic and environmental factors on AT2 cell function.

Our findings reveal intrinsic AT2 progenitor cell dysfunction in HPS, occurring early in the disease course prior to the onset of fibrosis and suggesting the potential for early targeted intervention. In addition, our findings reveal common pathways driving the fibrotic response across sporadic and genetic forms of pulmonary fibrosis. The relative contributions of rare and common genetic risk variants, in conjunction with environmental exposures, may vary across different forms of pulmonary fibrosis, but it is likely that these pathways converge, resulting in failure to maintain alveolar epithelial integrity.

Understanding the mechanisms that drive AT2 cell differentiation in HPS-PF and the optimal method to modulate the p53 pathway to promote epithelial maintenance and repair and prevent fibrotic remodeling will be critical. Additional studies at single-cell resolution may enable further characterization of the subpopulations of epithelial cells in HPS. We also found features of functional divergence between HPS1 and HPS2 AT2 cells in an organoid model and an influenza injury model, highlighting that the differences that exist between the HPS subtypes may help illuminate key aspects of HPS biology and have clinical relevance. Given the shared pathways that have been identified, further study of the role of alveolar epithelial dysfunction in HPS-PF should have broad applicability across genetic and sporadic fibrotic lung diseases.

## Methods

### Sex as a biological variable

For studies involving mice, both male and female mice were utilized, as fibrosis occurs equally in patients with HPS of both sexes.

### Animal studies and treatment

Naturally occurring HPS1, HPS2, and HPS3 mice, which harbor mutations in the mouse homologues of the *Hps1*, *Ap3b1* (*Hps2*), and *Hps3* genes, respectively, were maintained on the C57BL6/J background (47). Transgenic HPS2 (HPS2 TG) mice, with partial correction of AP3b1 in the lung epithelium driven by human-SFTPC promoter, as described previously (2). Double mutant HPS1/2 mice were generated by mating homozygous HPS1 and homozygous HPS2 mice (11, 13, 48). WT *Sftpc*^CreERT2/+^;*R26R*^EYFP/+^ (Jackson Laboratory, strain # 006148) mice were provided by Dr. David Frank and generated by Dr. Harold Chapman (University of California, San Francisco) as described previously, and bred with HPS1, HPS2, and HPS1/2 mice to generate HPS1 *Sftpc*^CreERT2/+^;*R26R*^EYFP/+^, HPS2 *Sftpc*^CreERT2/+^;*R26R*^EYFP/+^, and HPS1/2 *Sftpc*^CreERT2/+^;*R26R*^EYFP/+^ mice, respectively (49). Four- to 48-week-old animals were used for experiments, as specified. For lineage trace experiments, four-week-old *Sftpc*^CreERT2/+^;*R26R*^EYFP/+^ mice were administered tamoxifen (200 mg/kg body weight) in corn oil via oral gavage for three consecutive days. Keratinocyte growth factor (KGF) was obtained as palifermin (Amgen). KGF dissolved in vehicle (PBS) at a dose of 5 mg/kg or vehicle alone was administrated via intratracheal injection 48 hours prior to euthanasia and isolation of lungs (50). For cell proliferation experiments, pulse-chase 5-ethynyl-2’-deoxyuridine (EdU) dissolved in filtered PBS was administered via intraperitoneal injection at a dose of 50 mg/kg four hours prior to euthanasia and isolation of lungs.

### Human lung tissue

Normal lung samples were obtained through the University of Pennsylvania Lung Biology Institute’s Human Lung Tissue Bank. These samples were obtained from de-identified non-used lungs donated for organ transplantation through an established protocol (PROPEL, approved by University of Pennsylvania Institutional Review Board) with informed consent in accordance with institutional and NIH procedures, and consent was provided by next of kin or healthcare proxy (51). HPS-1 patient tissue samples were obtained through the University of Pennsylvania as part of the Prospective Registry of Outcomes in Patients Electing Lung Transplantation (PROPEL) and Vanderbilt University (51). The University of Pennsylvania and Vanderbilt University Institutional Review Boards (IRBs) approved this study, and written informed consent was obtained. All patient information was removed before use.

### Histology and immunofluorescence

Lungs were harvested at a constant pressure of 30 cm H_2_O and were fixed overnight at 4°C in 4% paraformaldehyde (PFA). For whole-mount imaging, lung tissue was washed in PBS, embedded in 4% low-melting agarose, and sectioned at 150 μm thickness. For paraffin embedding, lung tissue was washed in PBS, dehydrated in a graded series of ethanol washes, embedded in paraffin wax, and sectioned at 7 μm thickness. For human tissue, tissue was cut from the distal regions of the human lungs. 2 cm by 2 cm pieces of peripheral parenchyma were washed five times in cold PBS and then placed in 2% PFA overnight. The tissue was washed with PBS and dehydrated to 100% ethanol over the subsequent 12 to 24 hours, paraffin embedded and then sectioned at a thickness of 8 µm. Hematoxylin and eosin (H&E) and Masson’s trichrome staining were performed according to standard procedures. Immunofluorescence (IF) staining was performed with antibodies for prosurfactant protein-C (proSP-C) (Invitrogen, PA5-71680), NKX2.1 (Invitrogen, MA5-13961), GFP (Abcam, ab6673), LAMP3 (Synaptic Systems, 391 005), AGER (R&D Systems, MAB1179), KRT8 (DSHB), CLDN4 (Invitrogen, PA5-32354), LGALS3 (Invitrogen, 14-5301-82), KRT19 (Invitrogen, SA30-06), and HTII-280 (Terrace Biotech). EdU staining was performed with the Click-iT EdU cell proliferation kit according to the manufacturer’s instructions (Invitrogen). Terminal deoxynucleotidyl transferase dUTP nick end labeling (TUNEL) staining was performed using TUNEL assay apoptosis detection kit per the manufacturer’s instructions (Biotium).

Slides were mounted in Vectashield Antifade Mounting Medium (Vector Labs). Images were captured using a Leica confocal microscope. All image quantification was performed on serial confocal images from z-stack max intensity composites. For quantification, 10 images per sample were randomly selected via the 405 nm (DAPI) channel. Manual counting of DAPI nuclei, proSP-C^+^ cells, NKX2.1^+^ cells, and EYFP^+^ cells was performed in a blinded fashion.

### Alveolar epithelial type II cell isolation

Murine lung single cell suspensions were prepared by dispase digestion (52). Briefly, dispase (50 U/mL) was instilled in lungs followed by agarose to ensure digestion of distal airspaces and prevent incorporation of epithelial cells from distal airways. Lungs were incubated in a dispase solution for 45 minutes at room temperature, then DNase I (Stem Cell) was added, and lungs were teased apart with forceps. Serial filtration through 70-µm and 40-µm cell strainers was performed to create a single cell suspension, followed by ACK lysis to remove red blood cells. AT2 cells were subsequently isolated by fluorescence activated cell sorting (FACS) using antibodies for cell surface markers via negative selection for CD45, positive selection for EpCAM, and negative selection for β4 integrin (CD104) (Biolegend) (14, 49). Samples of the AT2 cell suspensions were fixed and stained for proSP-C, with confirmation of purity > 95%.

### Lung organoid assays

AT2 cells were isolated as described above. Live cell numbers were verified using a hemocytometer and Trypan Blue solution. In each technical replicate, 5,000 AT2 cells were combined with 50,000 primary outgrowth WT murine lung fibroblasts in 50% Matrigel (growth factor reduced, phenol-red free) (Corning) and in small airway growth media (Lonza) with the following additives: insulin, transferrin, bovine pituitary extract, retinoic acid, and gentamicin (all from Lonza), with cholera toxin (Sigma) and FBS (Corning). The ROCK inhibitor thiazovivin (Stem Cell) was included in the media for the first two days and media was changed every 2 to 3 days. Images were captured over the course of culture using a Leica Thunder microscope. Images were processed in ImageJ with thresholding and then binarized. All organoids within a well were used for quantification. Organoid number and size were quantified using the analyze particles function in ImageJ with a cutoff of 1500 μm^2^. Colony forming efficiency was calculated by dividing the organoid number by the number of input AT2 cells (5,000). Organoids were fixed and stained with antibodies for proSP-C (Invitrogen, PA5-71680) and AGER (Rage) (R&D Systems, MAB1179).

### Influenza infection

H1N1 PR8 influenza virus was generously provided by Dr. Andrew Vaughan (University of Pennsylvania) and titration by TCID_50_ for infectivity was performed in MDCK cells (53). For AT2 cell proliferation studies, we performed an initial set of experiments with WT and HPS2 mice, aged 8 to 12-weeks. Mice were anesthetized with 4% isoflurane in 100% O_2_ via an anesthesia vaporizer system and H1N1 PR8 influenza was delivered intratracheally at a dose of 100 U diluted in 50 μl PBS. In a subsequent experiment, HPS1 and WT mice, aged 8 to 12-weeks, were anesthetized with xylazine and ketamine via intraperitoneal injection, and H1N1 PR8 influenza was delivered intranasally at a dose of 100 U diluted in 50 μl PBS. For weight and oxygen saturation measurements, a subsequent cohort of WT, HPS1, and HPS2 mice, aged 8 to 12-weeks, were also anesthetized with xylazine and ketamine via intraperitoneal injection, and H1N1 PR8 influenza was delivered intranasally at a dose of 100 U diluted in 50 μl PBS. Mouse weights were tracked daily starting from the day of infection. Mice were shaved and pulse oximetry measurements were obtained using MouseOx Plus (Starr Life Sciences) every second or third day starting from the day of infection for the time points noted.

### Lung injury assessment

To define regions of lung injury after influenza infection, an unbiased image classification machine learning model was used on H&E-stained sections to cluster image pixels by staining intensity, similar to previously described methodology (15, 29). Three regions of injury were defined: severely injured, mildly injured, and unaffected.

### Bulk mRNA sequencing

Total RNA was extracted from freshly isolated AT2 cells using the RNeasy Micro Kit (Qiagen) and stored at -80°C. Frozen total RNA was sent to Genewiz/Azenta Life Sciences (NJ, USA) for bulk mRNA-sequencing, using their standard RNA-seq service. Reads in FASTQ format were aligned to the NCBI mouse genome (mm10/GRCm38) and quantified at the gene level using the nf-core/rnaseq v3.8.1 Nextflow pipeline (54, 55). Differential gene expression analysis was performed using DESeq2 and scanpy (56, 57). Aggregate gene set scores, including the REACTOME cellular senescence score, were calculated on a per-sample basis by selecting genes in the corresponding gene set and scoring them using the JASMINE scoring metric (58).

### β-galactosidase staining

Murine AT2 cells were isolated as described above. AT2 cells were deposited on glass slides by cytospin centrifugation at 500 rpm for 5 minutes. Fixation and staining were performed using a β-galactosidase staining kit per the manufacturer’s instructions at pH 6.0 (Cell Signaling). Cells were counterstained with DAPI to enable quantification. Human HPS-1 lung tissue was fresh frozen and sectioned via cryotome. Fixation and staining were performed using a β-galactosidase staining kit per the manufacturer’s instructions (Invitrogen, C10851). Sections were then counterstained for proSP-C (Abcam) and DAPI. The images were captured using a Keyence inverted fluorescence microscope.

### Alveolosphere differentiation assay

Alveolosphere culture was performed as previously described (22, 23). AT2 cells were isolated as described above. Live cell numbers were verified using a hemocytometer and Trypan Blue solution. In each technical replicate, 7,000 AT2 cells were plated in 50% Matrigel (growth factor reduced, phenol-red free) (Corning). Alveolospheres were cultured in alveolar maintenance media with IL-1β and the ROCK inhibitor Y-27632 (Sigma) for the first four days, alveolar differentiation media for the next four days, then transitioned to alveolar maintenance media for the remaining six days in culture. Media was changed every 2 to 3 days. Pifithrin-α (PFTα) was added to the alveolar differentiation media at a concentration of 30 μM (59, 60). Organoids were fixed and stained with antibodies for LAMP3 (Synaptic Systems, 391 005) and AGER (Rage) (R&D Systems, MAB1179).

### Quantitative polymerase chain reaction

Alveolospheres were dissociated from Matrigel using Cell Recovery Solution (Corning). RNA was isolated from alveolospheres using Direct-zol RNA Microprep Kits (Zymo). Complementary DNA (cDNA) was synthesized from total RNA using High-Capacity cDNA Reverse Transcription Kits (Applied Biosystems). Quantitative polymerase chain reaction (qPCR) was performed using the TaqMan system (Applied Biosystems). qPCR primers are listed in **Supplemental Table 1**. Relative gene expression was calculated using the ΔΔCt method (61).

### Picrosirius red staining

Staining for collagen was performed using picrosirius red staining kit per the manufacturer’s instructions (Polysciences Inc.). Digital morphometric measurements and collagen quantitation were performed as previously described (62). WT lung tissue was normalized to a collagen area of 3.0%.

### Statistics

Statistical analyses were performed with GraphPad Prism. Two-tailed unpaired Student’s *t* tests were used for comparison between groups for parametric data and Mann-Whitney U-test for non-parametric data.

Results are presented as mean ± SEM. *p* values < .05 were considered significant. For multiple comparisons, the Benjamini-Hochberg procedure was utilized with a false discovery rate of 0.05. Adjusted *p* values < .05 were considered significant.

### Study approval

All procedures for animal experiments were performed under the guidance of and in compliance with all ethical regulations of the Children’s Hospital of Philadelphia Institutional Animal Care and Use Committee (IACUC). Human HPS-1 specimens were obtained at the time of lung transplantation under established protocols at the University of Pennsylvania 458 (IRB protocol #813685) and Vanderbilt University (IRB protocol #171657). The normal human donor samples used were from de-identified, non-used lungs donated for research (PROPEL, IRB protocol #813685) with approval of the University of Pennsylvania and Vanderbilt University IRBs and in accordance with NIH procedures (51). Informed consent was provided by next of kin or health care proxy.

### Data availability

Mouse AT2 cell bulk RNA sequencing data are available in the NCBI’s public functional genomics data repository Gene Expression Omnibus (GEO accession number GSE235628). Values for all data points in graphs are reported in the Supporting Data Values file.

## Grant funding

J.Y.W. received support through an NIH Ruth L. Kirschstein National Research Service Award Institutional Research Training Grant (2-T32-HL-007586-36). J.J.G. receives support through a Parker B. Francis fellowship. E.C. receives support through an NIH Research Project Grant (R01HL155821). J.A.Z. receives support through an NIH Pathway to Independence Award (R00-HL141684), an NIH Maximizing Investigators’ Research Award (R35-GM146835), and an ATS/Hermansky-Pudlak Syndrome Network Research Grant. D.B.F receives support through an NIH Mentored Clinical Scientist Research Career Development Award (K08-HL140129). L.R.Y. receives support through an NIH Research Project Grant (R01-HL119503), an NIH Midcareer Investigator Award in Patient-Oriented Research (K24-HL143281), and the Rare Lung Diseases Frontier Program at the Children’s Hospital of Philadelphia.

## Author contributions

J.Y.W. designed and performed experiments, analyzed and interpreted the data, and wrote and edited the manuscript. S.N.M. analyzed and interpreted the data and edited the manuscript. S.S. designed and performed experiments and edited the manuscript. B.J.B. designed experiments and edited the manuscript. R.J. and J.J.G. performed experiments and reviewed the manuscript. S.M.L., M.C.B., and E.C. provided access to human samples and assisted with tissue processing. J.B.K. designed experiments and provided access to human samples and assisted with tissue processing. J.A.K., J.A.Z., and D.B.F. designed experiments, interpreted the data, and edited the manuscript. L.R.Y. designed the study, interpreted the data, and wrote and edited the manuscript.

## Acknowledgements

We thank the participating patients for providing human lung tissue specimens. We would also like to thank Ruby Pan, Mary Kate Goldy, and Heather Kelley for technical assistance and Dr. Andrew Vaughan, Dr. Luis Rodriguez, and Dr. Michael Beers for experimental guidance.

## Notes

### Competing Interest Statement

The authors have declared no competing interest.

### Summary of Updates

Additional lineage tracing experiments, ex vivo organoid experiments, and human HPS-1 patient experiments.

## References

1. Hengst M, et al. Hermansky-Pudlak syndrome type 2 manifests with fibrosing lung disease early in childhood. Orphanet journal of rare diseases. 2018;13(1):1–11.

2. Young LR, et al. The alveolar epithelium determines susceptibility to lung fibrosis in Hermansky-Pudlak syndrome. American journal of respiratory and critical care medicine. 2012;186(10):1014–24.

3. Lawson WE, et al. Endoplasmic reticulum stress in alveolar epithelial cells is prominent in IPF: Association with altered surfactant protein processing and herpesvirus infection. American Journal of Physiology-Lung Cellular and Molecular Physiology. 2008;294(6):L1119–26.

4. Armanios MY, et al. Telomerase mutations in families with idiopathic pulmonary fibrosis. N Engl J Med. 2007;356(13):1317–26.

5. Katzen J, et al. An SFTPC BRICHOS mutant links epithelial ER stress and spontaneous lung fibrosis. JCI insight. 2019;4(6)

6. Ahuja S, et al. MAP1LC3B overexpression protects against Hermansky-Pudlak syndrome type-1-induced defective autophagy in vitro. American Journal of Physiology-Lung Cellular and Molecular Physiology. 2016;310(6):L519–31.

7. Cuevas-Mora K, et al. Hermansky-Pudlak syndrome-2 alters mitochondrial homeostasis in the alveolar epithelium of the lung. Respiratory research. 2021;22(1):1–11.

8. Zhou Y, et al. Chitinase 3–like–1 and its receptors in Hermansky-Pudlak syndrome–associated lung disease. J Clin Invest. 2015;125(8):3178–92.

9. Cinar R, et al. CB1R and iNOS are distinct players promoting pulmonary fibrosis in Hermansky– Pudlak syndrome. Clinical and Translational Medicine. 2021;11(7):e471.

10. Young LR, et al. Susceptibility of Hermansky-Pudlak mice to bleomycin-induced type II cell apoptosis and fibrosis. American journal of respiratory cell and molecular biology. 2007;37(1):67–74.

11. Mahavadi P, et al. Epithelial stress and apoptosis underlie Hermansky-Pudlak syndrome–associated interstitial pneumonia. American journal of respiratory and critical care medicine. 2010;182(2):207–19.

12. Atochina-Vasserman EN, et al. Early alveolar epithelial dysfunction promotes lung inflammation in a mouse model of Hermansky-Pudlak syndrome. American journal of respiratory and critical care medicine. 2011;184(4):449–58.

13. Lyerla TA, et al. Aberrant lung structure, composition, and function in a murine model of Hermansky-Pudlak syndrome. American Journal of Physiology-Lung Cellular and Molecular Physiology. 2003;285(3):L643–53.

14. Vaughan AE, et al. Lineage-negative progenitors mobilize to regenerate lung epithelium after major injury. Nature. 2015;517(7536):621-5.

15. Zacharias WJ, et al. Regeneration of the lung alveolus by an evolutionarily conserved epithelial progenitor. Nature. 2018;555(7695):251-5.

16. Strunz M, et al. Alveolar regeneration through a Krt8 transitional stem cell state that persists in human lung fibrosis. Nature communications. 2020;11(1):1–20.

17. Kobayashi Y, et al. Persistence of a regeneration-associated, transitional alveolar epithelial cell state in pulmonary fibrosis. Nat Cell Biol. 2020;22(8):934–46.

18. Choi J, et al. Inflammatory signals induce AT2 cell-derived damage-associated transient progenitors that mediate alveolar regeneration. Cell stem cell. 2020;27(3):366,382. e7.

19. Yao C, et al. Senescence of alveolar type 2 cells drives progressive pulmonary fibrosis. American Journal of Respiratory and Critical Care Medicine. 2021;203(6):707–17.

20. Parimon T, et al. Senescence of alveolar epithelial progenitor cells: A critical driver of lung fibrosis. American Journal of Physiology-Cell Physiology. 2023;325(2):C483–95.

21. Hamsanathan S, et al. Cellular senescence: The trojan horse in chronic lung diseases. American journal of respiratory cell and molecular biology. 2019;61(1):21–30.

22. Katsura H, et al. Human lung stem cell-based alveolospheres provide insights into SARS-CoV-2-mediated interferon responses and pneumocyte dysfunction. Cell stem cell. 2020;27(6):890,904. e8.

23. Konishi S, et al. Defined conditions for long-term expansion of murine and human alveolar epithelial stem cells in three-dimensional cultures. STAR protocols. 2022;3(2):101447.

24. Parimon T, et al. Alveolar epithelial type II cells as drivers of lung fibrosis in idiopathic pulmonary fibrosis. International journal of molecular sciences. 2020;21(7):2269.

25. Katzen J, Beers MF. Contributions of alveolar epithelial cell quality control to pulmonary fibrosis. J Clin Invest. 2020;130(10):5088–99.

26. Barkauskas CE, et al. Type 2 alveolar cells are stem cells in adult lung. J Clin Invest. 2013;123(7):3025–36.

27. Zhang D, et al. Rare and common variants in KIF15 contribute to genetic risk of idiopathic pulmonary fibrosis. American journal of respiratory and critical care medicine. 2022;206(1):56–69.

28. Young LR, Borchers MT, Allen HL, Gibbons RS, McCormack FX. Lung-restricted macrophage activation in the pearl mouse model of hermansky-pudlak syndrome. The Journal of Immunology. 2006;176(7):4361–8.

29. Liberti DC, et al. Alveolar epithelial cell fate is maintained in a spatially restricted manner to promote lung regeneration after acute injury. Cell reports. 2021;35(6):109092.

30. Nabhan AN, et al. Single-cell Wnt signaling niches maintain stemness of alveolar type 2 cells. Science. 2018;359(6380):1118-23.

31. Jansing NL, et al. Unbiased quantitation of alveolar type II to alveolar type I cell transdifferentiation during repair after lung injury in mice. American journal of respiratory cell and molecular biology. 2017;57(5):519–26.

32. Wang F, et al. Regulation of epithelial transitional states in murine and human pulmonary fibrosis. J Clin Invest. 2023;133(22)

33. Konkimalla A, et al. Transitional cell states sculpt tissue topology during lung regeneration. Cell Stem Cell. 2023;30(11):1486,1502. e9.

34. Katzen J, et al. Disruption of proteostasis causes IRE1 mediated reprogramming of alveolar epithelial cells. Proceedings of the National Academy of Sciences. 2022;119(43):e2123187119.

35. Toth A, et al. Alveolar epithelial progenitor cells require Nkx2-1 to maintain progenitor-specific epigenomic state during lung homeostasis and regeneration. Nature Communications. 2023;14(1):8452.

36. Alder JK, et al. Telomere dysfunction causes alveolar stem cell failure. Proceedings of the National Academy of Sciences. 2015;112(16):5099–104.

37. Young LR, et al. Epithelial-macrophage interactions determine pulmonary fibrosis susceptibility in Hermansky-Pudlak syndrome. JCI insight. 2016;1(17)

38. Rouhani FN, et al. Alveolar macrophage dysregulation in Hermansky-Pudlak syndrome type 1. American journal of respiratory and critical care medicine. 2009;180(11):1114–21.

39. Flaherty KR, et al. Histopathologic variability in usual and nonspecific interstitial pneumonias. American journal of respiratory and critical care medicine. 2001;164(9):1722–7.

40. Korfei M, et al. Epithelial endoplasmic reticulum stress and apoptosis in sporadic idiopathic pulmonary fibrosis. American journal of respiratory and critical care medicine. 2008;178(8):838–46.

41. Thannickal VJ, Horowitz JC. Evolving concepts of apoptosis in idiopathic pulmonary fibrosis. Proceedings of the American Thoracic Society. 2006;3(4):350–6.

42. Riemondy KA, Jansing NL, Jiang P, Redente EF, Gillen AE, Fu R, Miller AJ, Spence JR, Gerber AN, Hesselberth JR. Single-cell RNA sequencing identifies TGF-β as a key regenerative cue following LPS-induced lung injury. JCI Insight. 2019;4(8).

43. Joshi N, Watanabe S, Verma R, Jablonski RP, Chen C, Cheresh P, Markov NS, Reyfman PA, McQuattie-Pimentel AC, Sichizya L. A spatially restricted fibrotic niche in pulmonary fibrosis is sustained by M-CSF/M-CSFR signalling in monocyte-derived alveolar macrophages. European Respiratory Journal. 2020;55(1).

44. Wu H, Yu Y, Huang H, Hu Y, Fu S, Wang Z, Shi M, Zhao X, Yuan J, Li J. Progressive pulmonary fibrosis is caused by elevated mechanical tension on alveolar stem cells. Cell. 2020;180(1):107,121. e17.

45. Watanabe S, Markov NS, Lu Z, Piseaux Aillon R, Soberanes S, Runyan CE, Ren Z, Grant RA, Maciel M, Abdala-Valencia H. Resetting proteostasis with ISRIB promotes epithelial differentiation to attenuate pulmonary fibrosis. Proceedings of the National Academy of Sciences. 2021;118(20):e2101100118.

46. Auyeung VC, Downey MS, Thamsen M, Wenger TA, Backes BJ, Sheppard D, Papa FR. IRE1α drives lung epithelial progenitor dysfunction to establish a niche for pulmonary fibrosis. American Journal of Physiology Lung Cellular and Molecular Physiology. 2022;322(4):L564–L580.

47. Li W, et al. Murine Hermansky–Pudlak syndrome genes: Regulators of lysosome-related organelles. Bioessays. 2004;26(6):616–28.

48. Guttentag SH, et al. Defective surfactant secretion in a mouse model of Hermansky-Pudlak syndrome. American journal of respiratory cell and molecular biology. 2005;33(1):14–21.

49. Chapman HA, et al. Integrin α6β4 identifies an adult distal lung epithelial population with regenerative potential in mice. J Clin Invest. 2011;121(7):2855–62.

50. Yano T, et al. KGF regulates pulmonary epithelial proliferation and surfactant protein gene expression in adult rat lung. American Journal of Physiology-Lung Cellular and Molecular Physiology. 2000;279(6):L1146–58.

51. Diamond JM, et al. Clinical risk factors for primary graft dysfunction after lung transplantation. American journal of respiratory and critical care medicine. 2013;187(5):527–34.

52. Sinha M, Lowell CA. Isolation of highly pure primary mouse alveolar epithelial type II cells by flow cytometric cell sorting. Bio-protocol. 2016;6(22):e2013.

53. Weiner AI, et al. ΔNp63 drives dysplastic alveolar remodeling and restricts epithelial plasticity upon severe lung injury. Cell reports. 2022;41(11):111805.

54. Di Tommaso P, et al. Nextflow enables reproducible computational workflows. Nat Biotechnol. 2017;35(4):316–9.

55. Ewels PA, et al. The nf-core framework for community-curated bioinformatics pipelines. Nat Biotechnol. 2020;38(3):276–8.

56. Love MI, et al. Moderated estimation of fold change and dispersion for RNA-seq data with DESeq2. Genome Biol. 2014;15(12):1–21.

57. Wolf FA, et al. SCANPY: Large-scale single-cell gene expression data analysis. Genome Biol. 2018;19:1–5.

58. Noureen N, et al. Signature-scoring methods developed for bulk samples are not adequate for cancer single-cell RNA sequencing data. Elife. 2022;11:e71994.

59. Zhu J, et al. Pifithrin-α alters p53 post-translational modifications pattern and differentially inhibits p53 target genes. Scientific reports. 2020;10(1):1049.

60. Sohn D, et al. Pifithrin-α protects against DNA damage-induced apoptosis downstream of mitochondria independent of p53. Cell Death & Differentiation. 2009;16(6):869–78.

61. Livak KJ, Schmittgen TD. Analysis of relative gene expression data using real-time quantitative PCR and the 2− ΔΔCT method. Methods. 2001;25(4):402–8.

62. Nureki S, et al. Expression of mutant Sftpc in murine alveolar epithelia drives spontaneous lung fibrosis. J Clin Invest. 2018;128(9):4008–24.

