## Supplemental Figures and Tables for "Dysregulated alveolar epithelial cell progenitor function and identity in Hermansky-Pudlak syndrome"

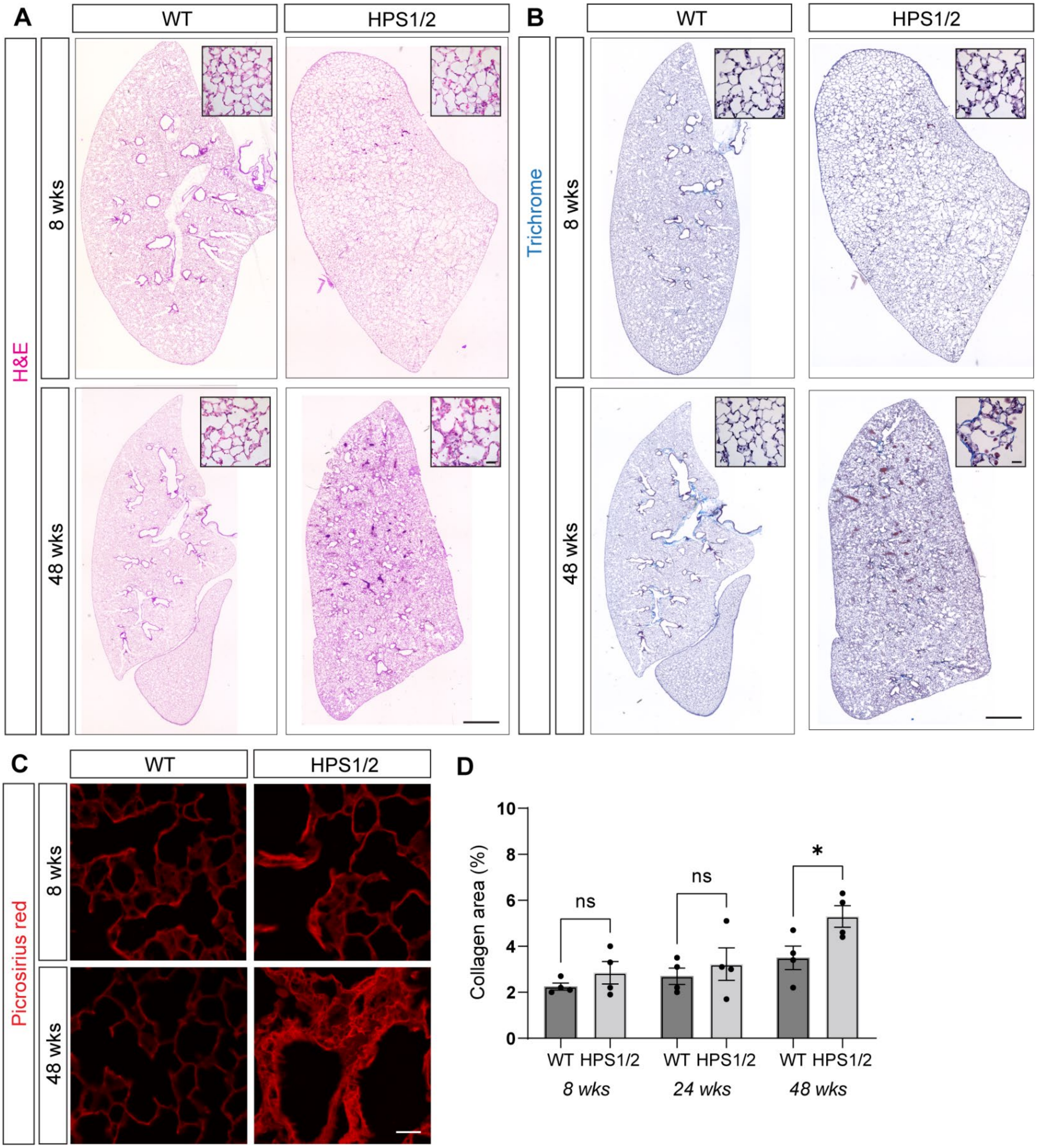


**Supplemental Figure 1. Progressive pulmonary fibrosis with aging in HPS1/2 mice.** (A) H&E and (B) trichrome stained whole lung section images with insets and (C) picrosirius red stained images from WT and HPS1/2 mice at 8 and 48 weeks of age. (D) Quantification of the percentage of collagen area detected by picrosirius red staining in WT and HPS1/2 mice at 8, 24, and 48 weeks of age. All quantification data are represented as mean ± SEM. Two-tailed t tests: ns, not significant; * $p < .05$; n = 4 per group per time point. Scale bars in (A, B) 1 mm with 20 μm inset; (C) 20 μm.

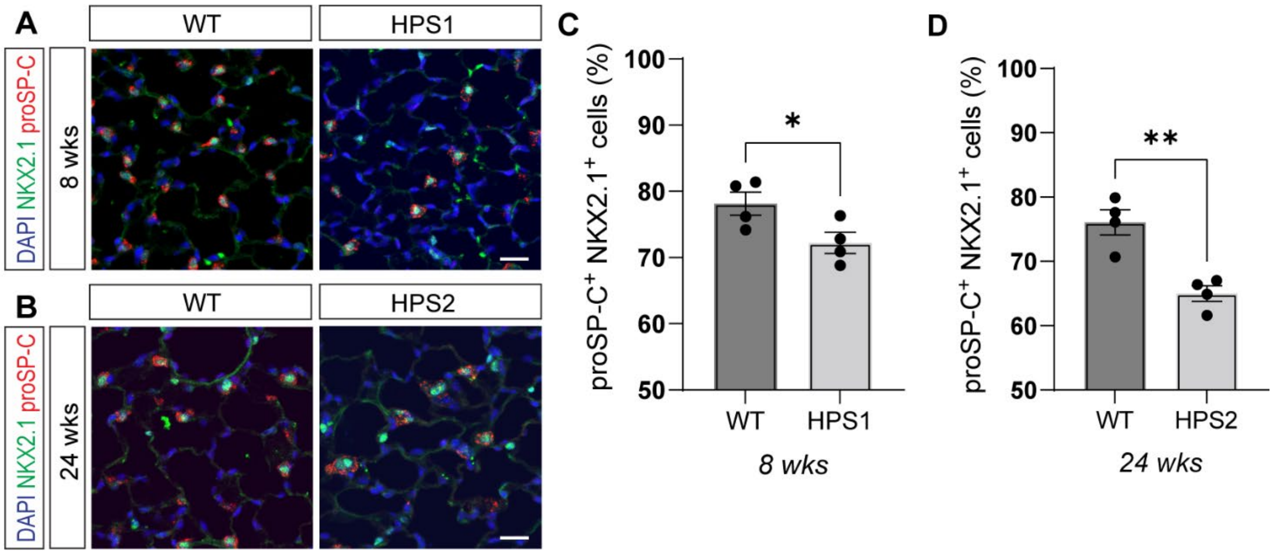


**Supplemental Figure 2. Quantification of alveolar type II epithelial (AT2) cells as a percentage of total epithelial cells in HPS1 and HPS2 mice.** (A-B) Immunofluorescence staining of paraffin-embedded lung tissue for proSP-C and NKX2.1 in (A) WT and HPS1 mice at 8 weeks of age and (B) WT and HPS2 mice at 24 weeks of age. (C-D) Quantification of percentage of proSP-$C^+$ NKX2.$1^+$ cells as a percentage of total epithelial cells (by NKX2.$1^+$ cells) in (C) WT vs. HPS1 mice at 8 weeks of age and (D) WT vs. HPS2 mice at 24 weeks of age. DAPI stains nuclei (blue). All quantification data are represented as mean ± SEM. Two-tailed t tests: * $p < .05$, ** $p < .01$; n = 4 per group per time point. Scale bars in (A, B), 20 µm.

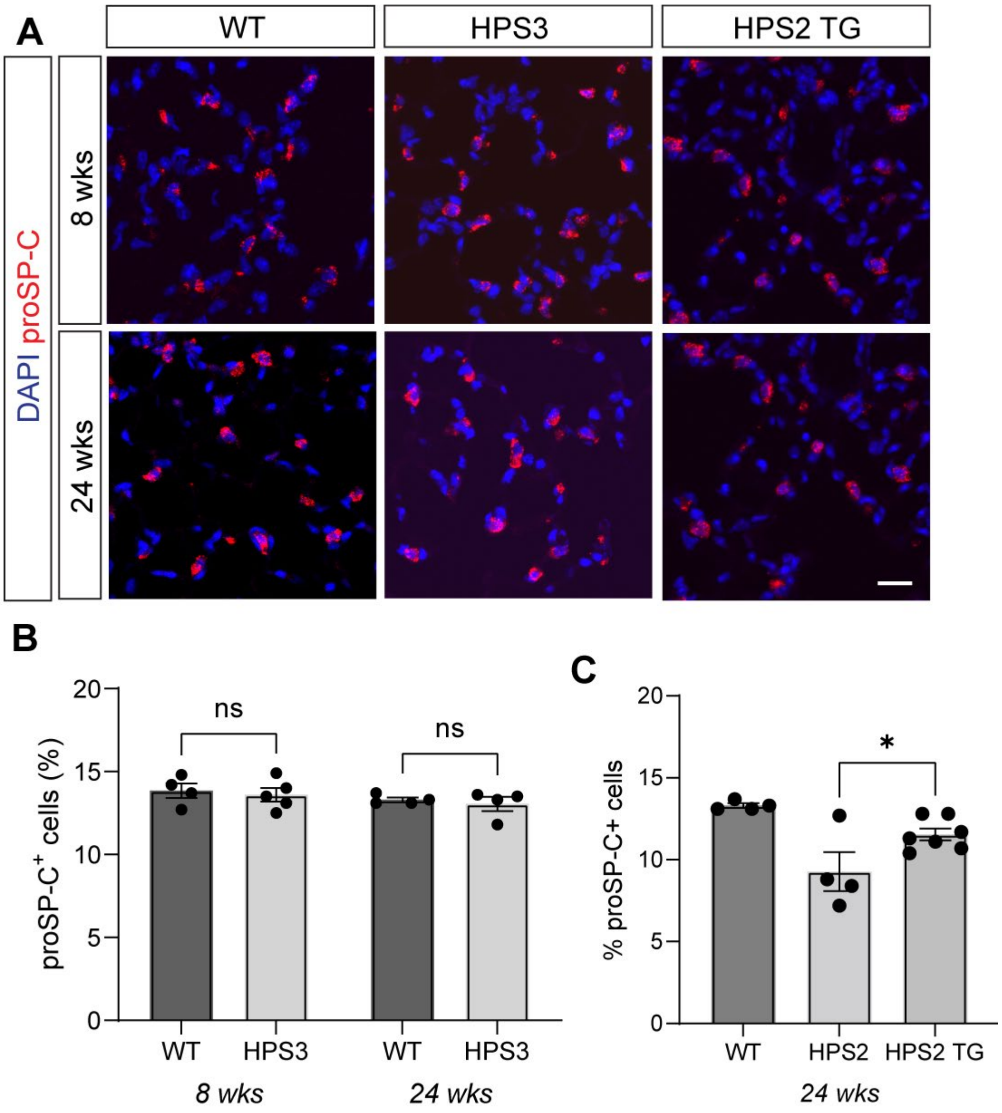


**Supplemental Figure 3. Quantification of alveolar type II epithelial (AT2) cells with aging in HPS3 and HPS2 transgenic epithelial-corrected (HPS2 TG) mice.** (A) Immunofluorescence staining of paraffin-embedded lung tissue for proSP-C in WT, HPS3, and HPS2 transgenic epithelial-corrected (HPS2 TG) mice at 8 and 24 weeks of age. (B, C) Quantification of percentage of proSP-$C^+$ cells as a percentage of total cells (by DAPI staining) in (B) WT vs. HPS3 mice at 8 and 24 weeks of age and (C) WT vs. HPS2 TG mice at 24 weeks of age. DAPI stains nuclei (blue). All quantification data are represented as mean ± SEM. Two-tailed t tests: ns, not significant; * $p < .05$; n = 4-7 per group per time point. Scale bars in (A), 20 µm.

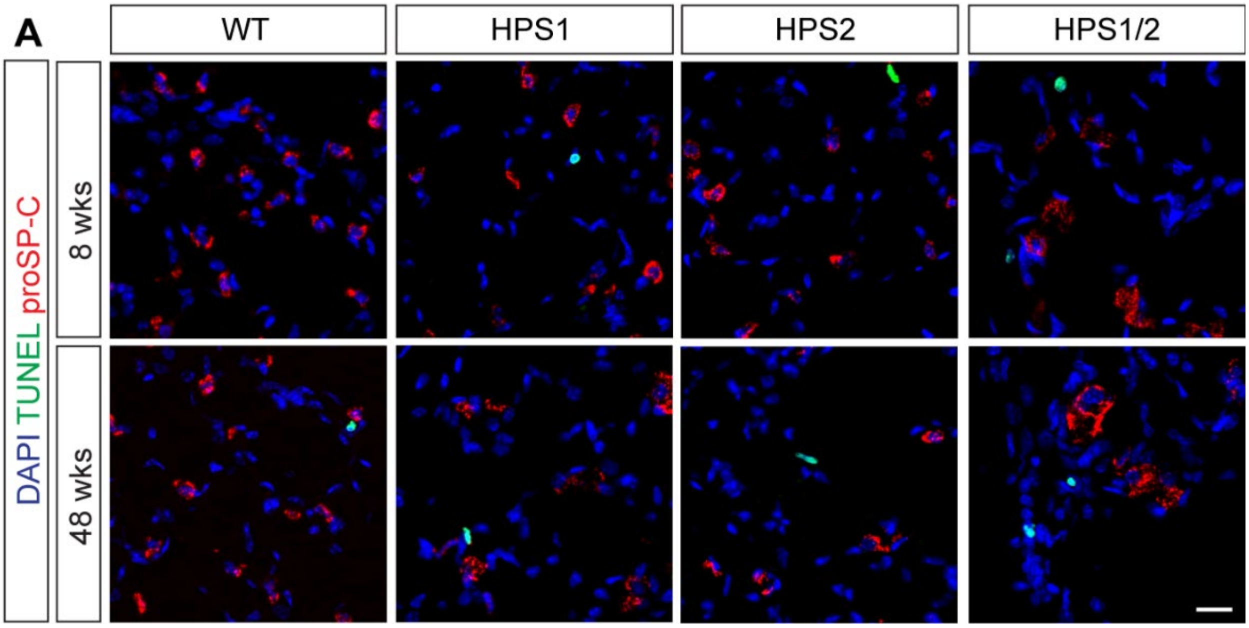


**Supplemental Figure 4. Evaluation for alveolar type II epithelial (AT2) cell apoptosis with aging in HPS mice.** (A) Immunofluorescence staining of paraffin-embedded lung tissue for TUNEL and proSP-C in WT, HPS1, HPS2, and HPS1/2 mice at 8 and 48 weeks of age. DAPI stains nuclei (blue). Scale bars in (A), 20 µm.

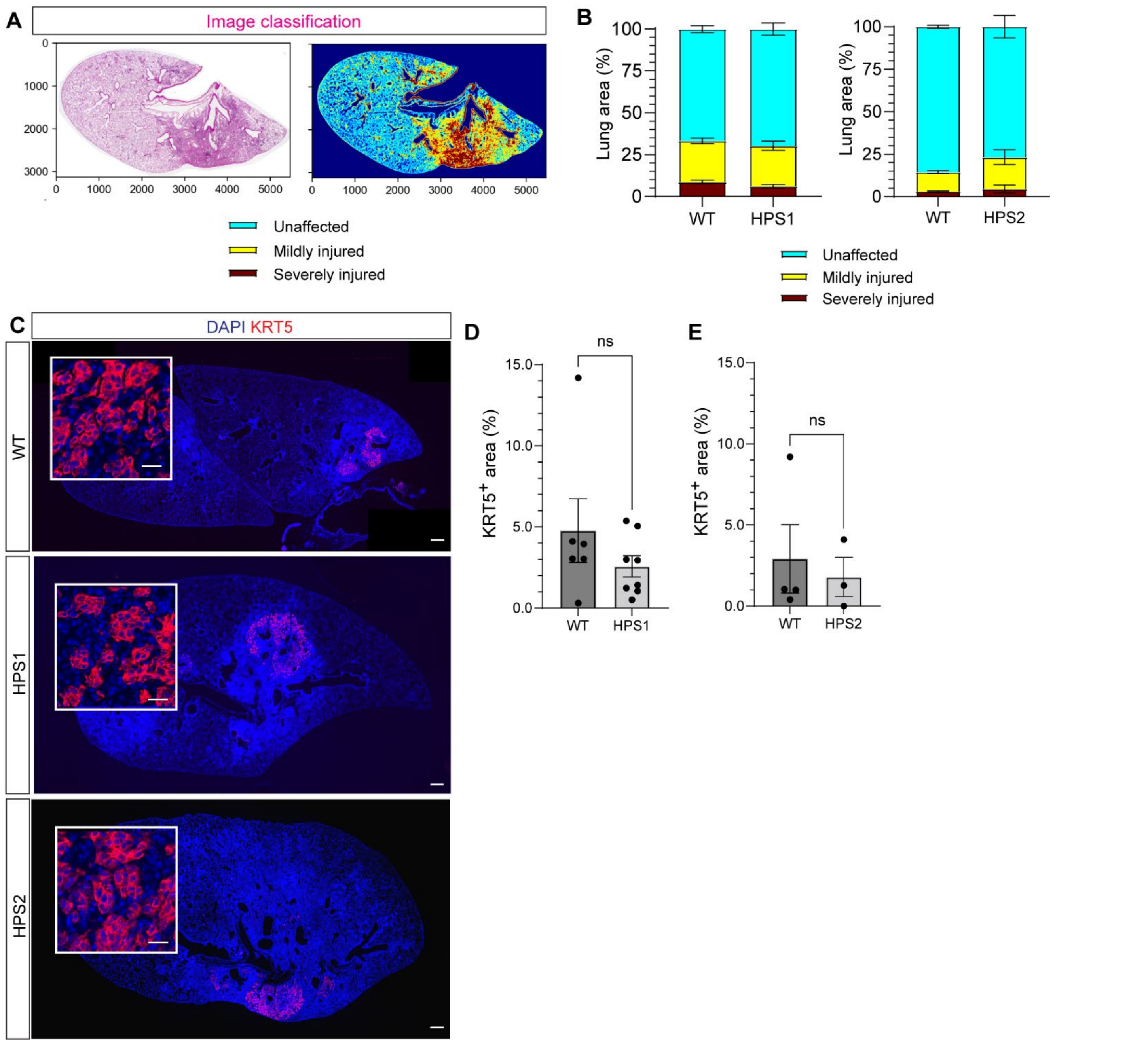


**Supplemental Figure 5. Injury severity in acute influenza infection in HPS mice.** (A) Computational image analysis to define regions of lung injury from H&E-stained lung sections. (B) Percentage of unaffected, mildly injured, and severely injured regions in WT vs. HPS1 mice and WT. vs. HPS2 mice at 14 days post-infection (dpi) with influenza. (C) Immunofluorescence staining of lung tissue for keratin 5 (KRT5) in WT, HPS1, and HPS2 mice at 14 dpi. (D, E) Quantification of $KRT5^+$ area as a percentage of total lung area in (D) WT vs. HPS1 mice and (E) WT vs. HPS2 mice at 14 dpi. DAPI stains nuclei (blue). All quantification data are represented as mean ± SEM. Two-tailed t tests: ns, not significant; n = 3-8 per group. Scale bars in (A), 2 cm; inset, 20 µm.

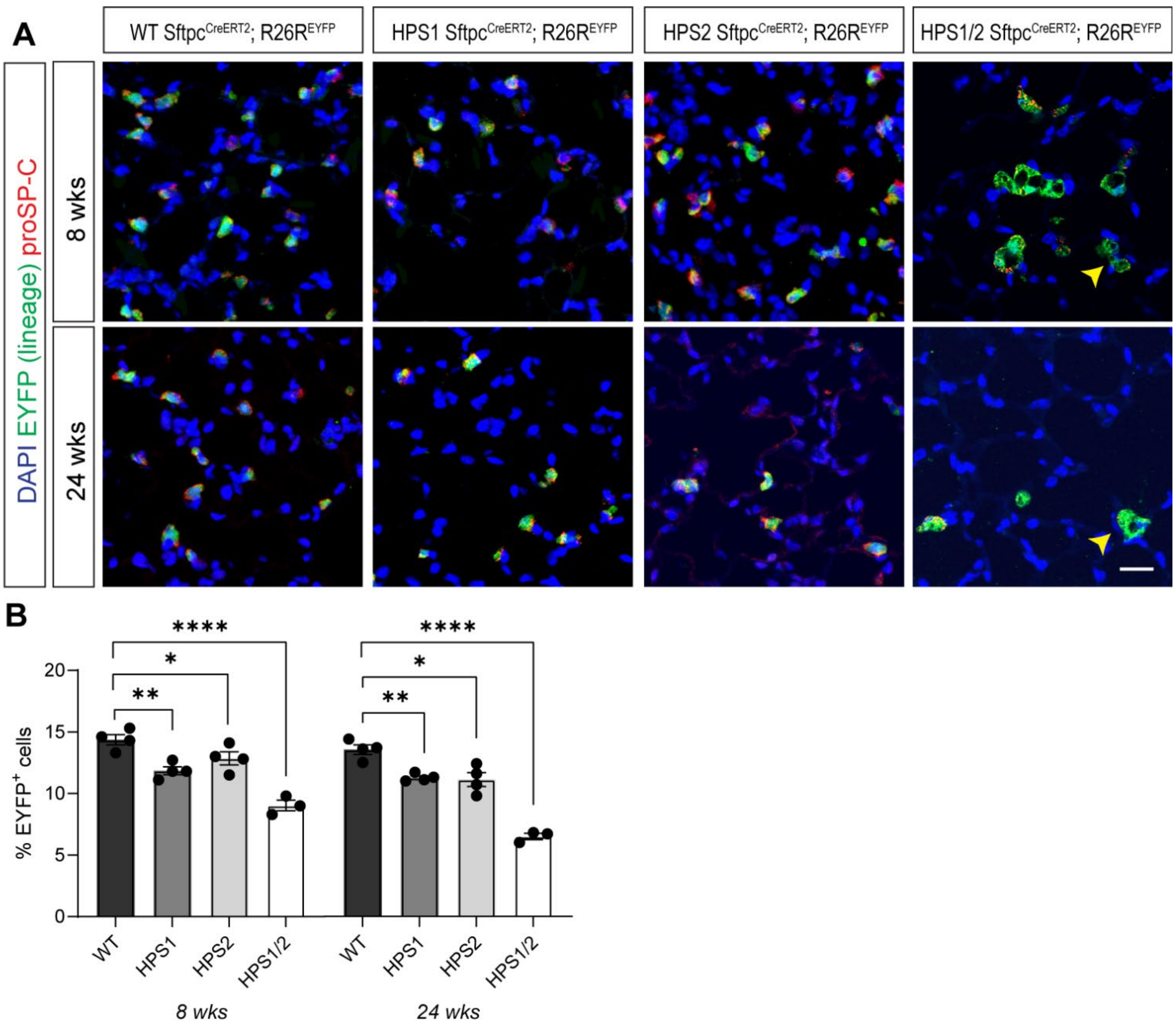


**Supplemental Figure 6. Lineage tracing of alveolar type II epithelial (AT2) cells in HPS mice confirms loss of AT2 cells.** (A) Immunofluorescence staining of paraffin-embedded lung tissue for proSP-C and the lineage marker EYFP in WT, HPS1, HPS2, and HPS1/2 Sftpc$^{CreERT2/+}$;R26R$^{EYFP/+}$ mice at 8 and 24 weeks of age. Arrows indicate EYFP$^{+}$, proSP-C$^{neg}$ cells. (B) Quantification of EYFP$^{+}$ cells as a percentage of total cells (by DAPI staining). DAPI stains nuclei (blue). All quantification data are represented as mean ± SEM. Statistics using two-tailed unpaired Student's *t* tests: adjusted * $p < .05$; ** $p < .01$; **** $p < .0001$ after Benjamini Hochberg correction for multiple comparisons. n = 3-4 per group per time point. Scale bars in (A), 20 μm.

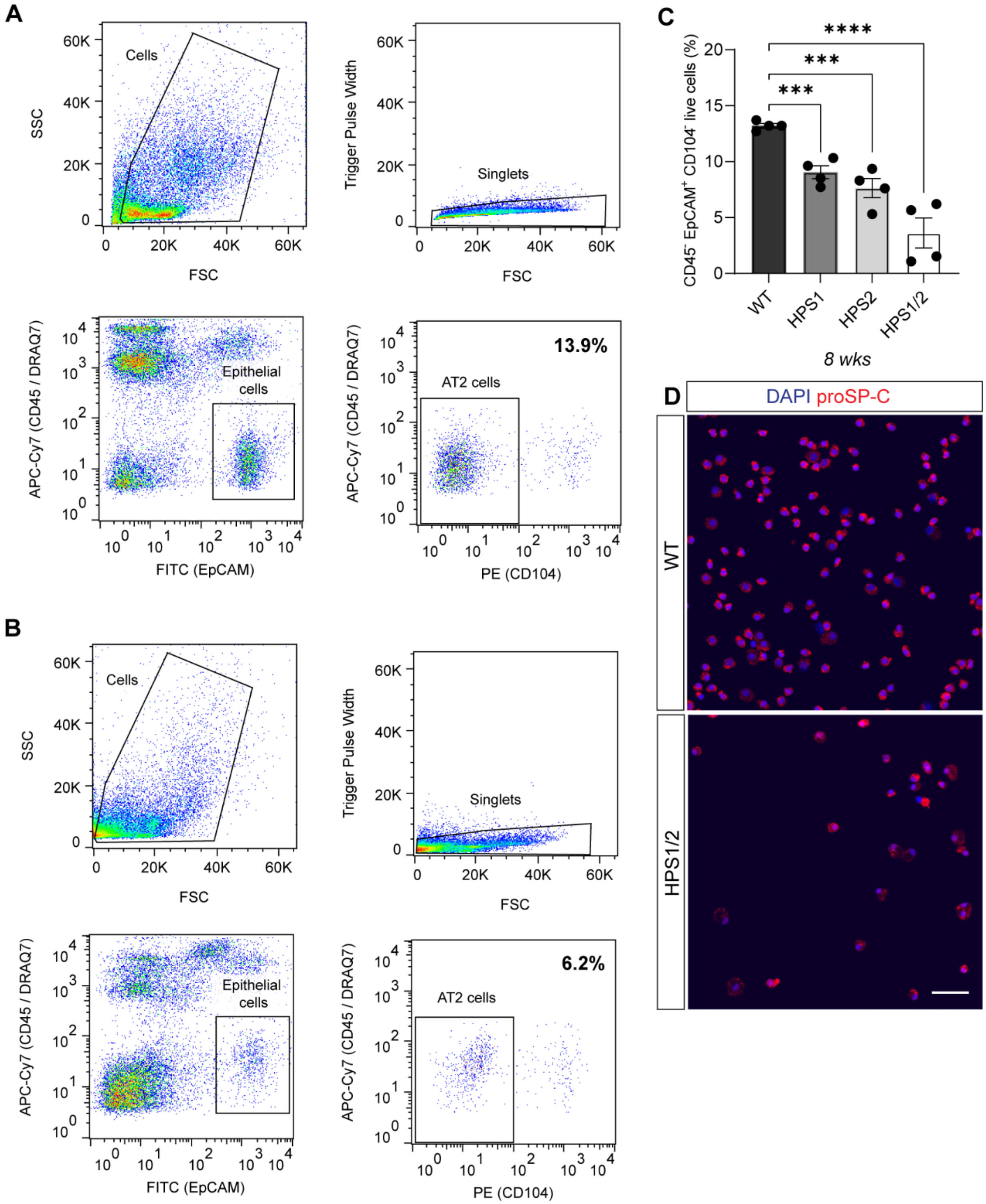


**Supplemental Figure 7. Reduced AT2 cell percentages in HPS mice by fluorescence activated cell sorting (FACS).** (A-B) Representative FACS plots of $CD45^-$ $EpCAM^+$ $CD104^-$ live AT2 cell population in (A) WT and (B) HPS1/2 mice at 8 weeks of age. (C) Quantification of percentage of $CD45^-$ $EpCAM^+$ $CD104^-$ live AT2 cells in WT, HPS1, HPS2, and HPS1/2 mice at 8 weeks of age. (D) proSP-C immunostaining of cytospin preparations from AT2 cells isolated from WT and HPS1/2 mice at 8 weeks of age. DAPI stains nuclei (blue). All quantification data are represented as mean ± SEM. Statistics using two-tailed unpaired Student's *t* tests: adjusted *** $p < .001$; **** $p < .0001$ after Benjamini Hochberg correction for multiple comparisons. n = 4 per group per time point. Scale bars in (D), 50 µm.

**A**

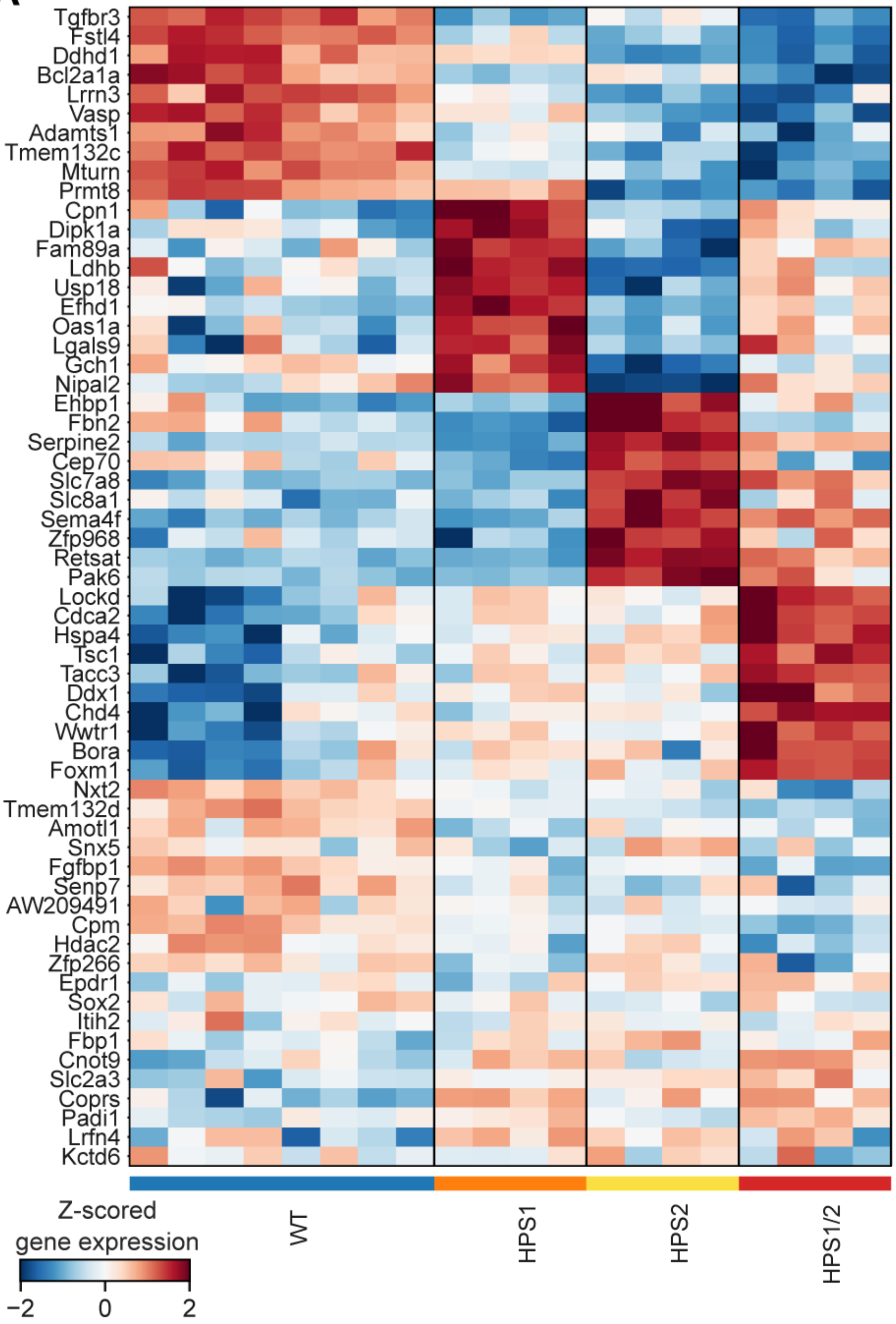
Tgfbr3
Fstl4
Ddhd1
Bcl2a1a
Lrrn3
Vasp
Adamts1
Tmem132c
Mturn
Prmt8
Cpn1
Dipk1a
Fam89a
Ldhb
Usp18
Efhd1
Oas1a
Lgals9
Gch1
Nipal2
Ehbp1
Fbn2
Serpine2
Cep70
Slc7a8
Slc8a1
Sema4f
Zfp968
Retsat
Pak6
Lockd
Cdca2
Hspa4
Tsc1
Tacc3
Ddx1
Chd4
Wwtr1
Bora
Foxm1
Nxt2
Tmem132d
Amotl1
Snx5
Fgfbp1
Senp7
AW209491
Cpm
Hdac2
Zfp266
Epdr1
Sox2
Itih2
Fbp1
Cnot9
Slc2a3
Coprs
Padi1
Lrfn4
Kctd6
Z-scored gene expression
−2
0
2
WT
HPS1
HPS2
HPS1/2

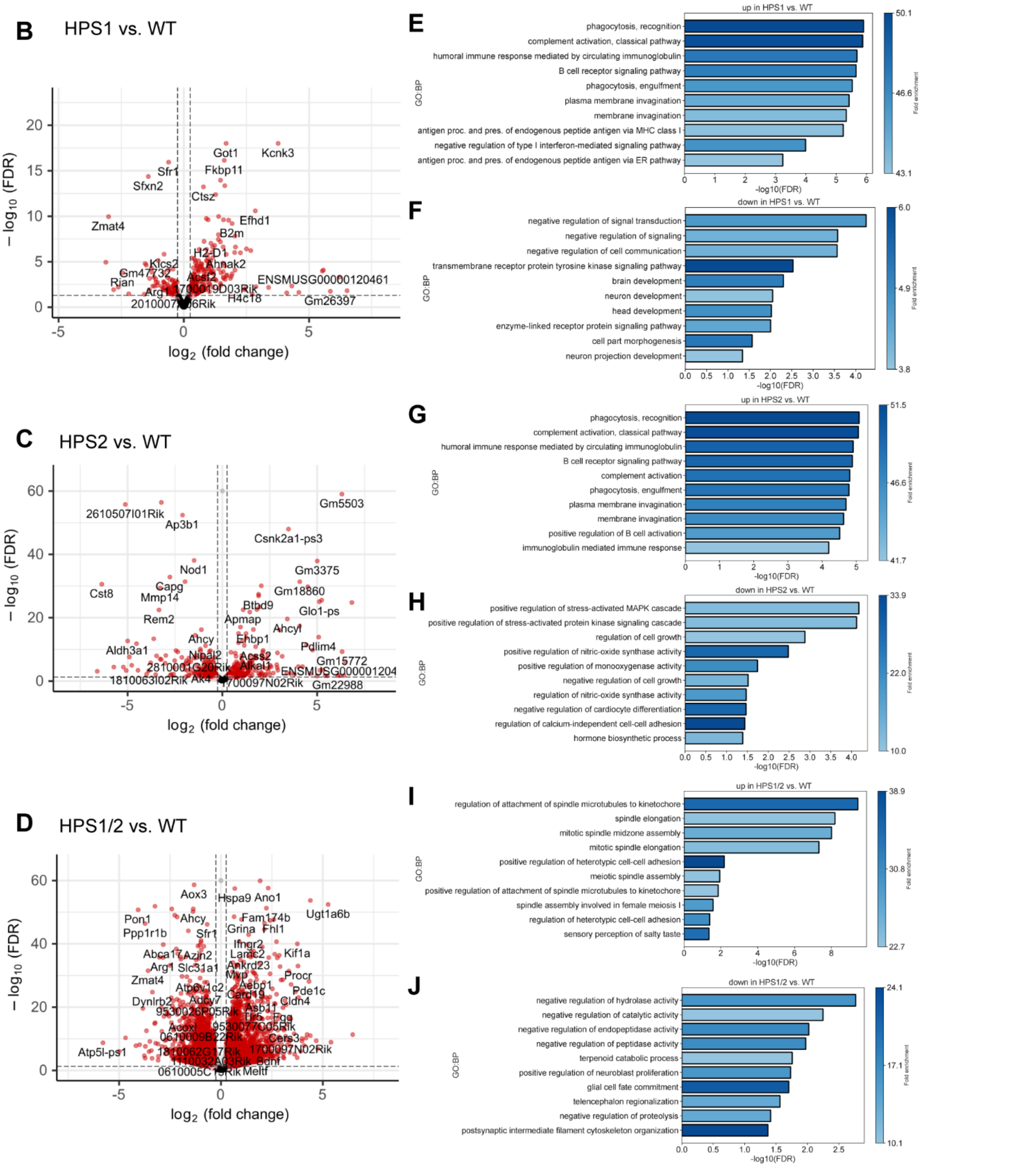


**Supplemental Figure 8. Differential gene expression analysis of WT vs. HPS AT2 cells.** (A) Matrixplot of top genes over- and under-expressed in AT2 cells from WT, HPS1, HPS2, and HPS1/2 mice at 8 weeks of age. (B-D) Volcano plots showing the log2-fold changes (LFCs) in gene expression as a function of genotype in (B) HPS1, (C) HPS2, and (D) HPS1/2 vs. WT AT2 cells. (E-J) Over-representation analysis (ORA) from gProfiler showing GO: Biological Processes (GO:BP) associated with genes upregulated in (E) HPS1, (G) HPS2, and (I) HPS1/2 vs. WT AT2 cells and downregulated in (F) HPS1, (H) HPS2, and (J) HPS1/2 vs. WT AT2 cells.

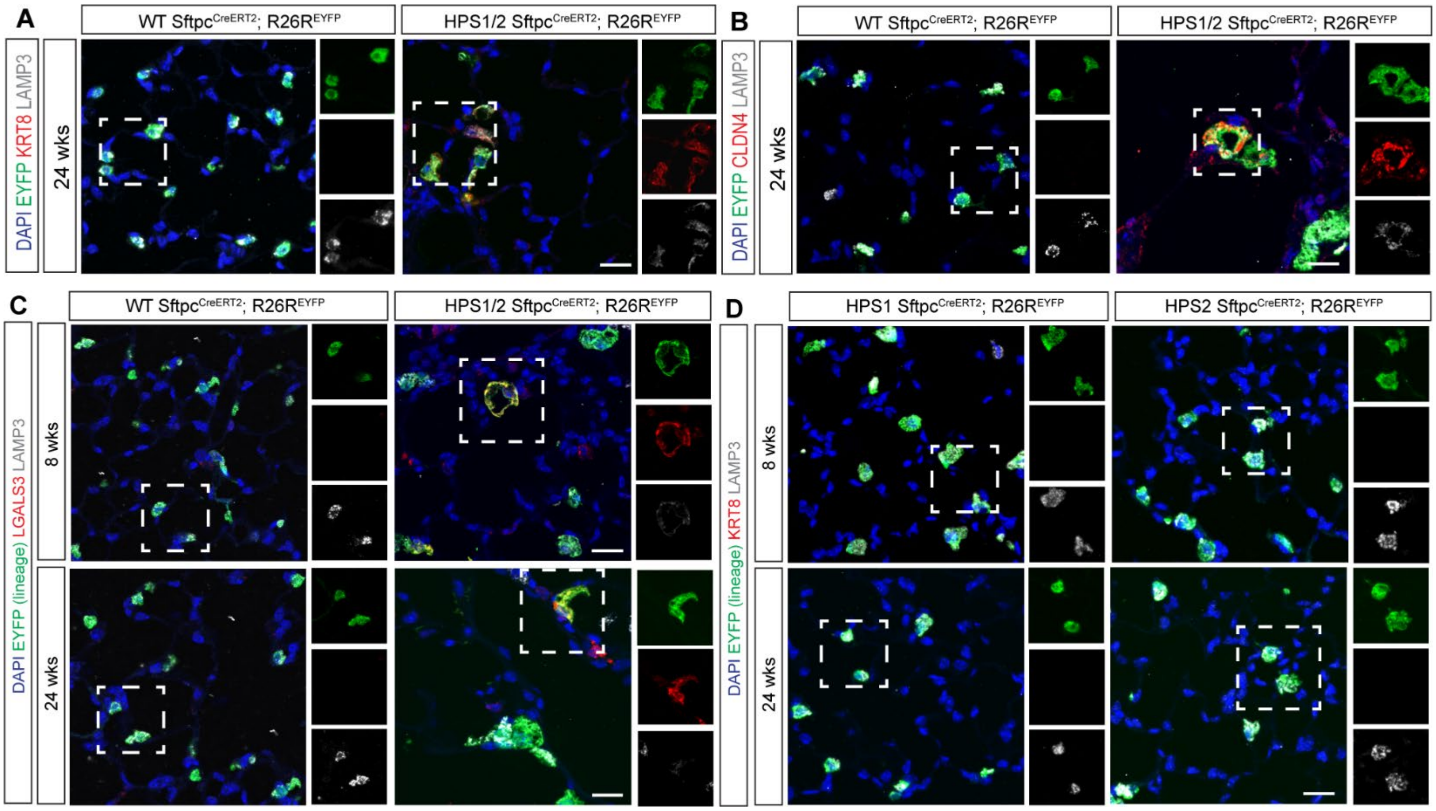


**Supplemental Figure 9. Evaluation for reprogrammed transitional cells in HPS mice.** (A, B) Immunofluorescence (IF) staining of paraffin-embedded lung tissue for EYFP, (A) KRT8 or (B) CLDN4, and LAMP3 in WT and HPS1/2 Sftpc$^{CreERT2/+}$;R26R$^{EYFP/+}$ mice at 24 weeks of age. (C) IF staining for EYFP, LGALS3, and LAMP3 in WT and HPS1/2 Sftpc$^{CreERT2/+}$;R26R$^{EYFP/+}$ mice at 8 and 24 weeks of age. (D) IF staining for EYFP, KRT8, and LAMP3 in HPS1 and HPS2 Sftpc$^{CreERT2/+}$;R26R$^{EYFP/+}$ mice at 8 and 24 weeks of age. Dashed line boxes represent EYFP$^{+}$ with inset boxes displaying EYFP; KRT8, CLDN4 or KRT8; and LAMP3 staining. DAPI stains nuclei (blue). All scale bars, 20 µm.

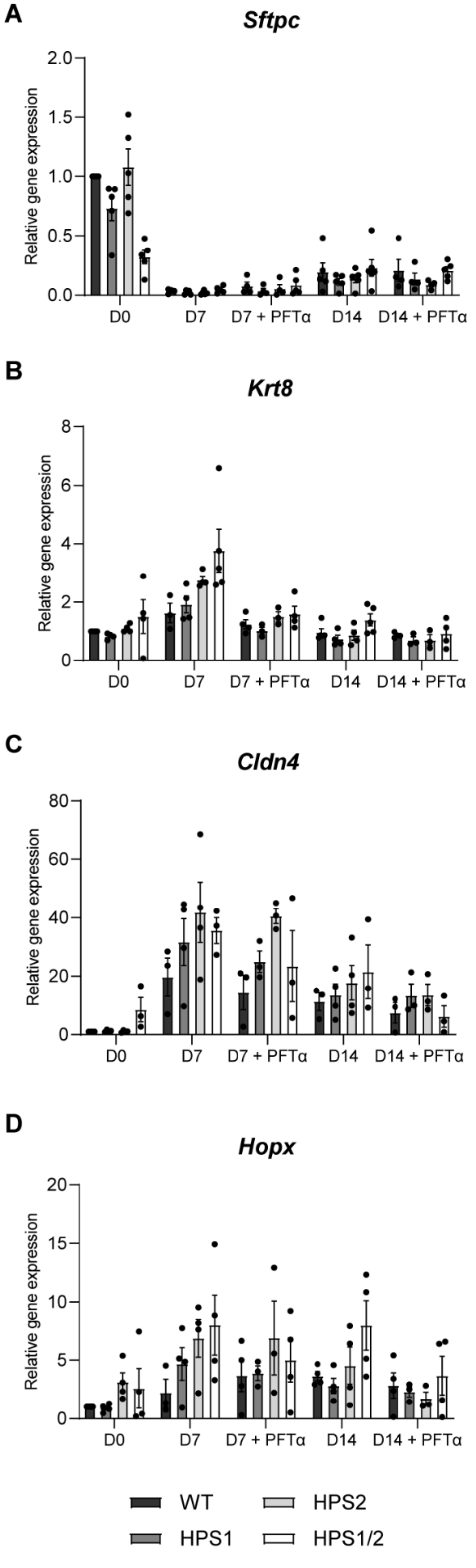


**Supplemental Figure 10. Gene expression of WT, HPS1, HPS2, and HPS1/2 alveolospheres.** (A-D) Relative gene expression of selected (A) alveolar epithelial type II (AT2) cell (*Sftpc*), (B, C) reprogrammed transitional cell (*Krt8*, *Cldn4*), and (D) alveolar epithelial type I (AT1) cell (*Hopx*) genes in alveolospheres generated from AT2 cells from 8-week-old WT, HPS1, HPS2, and HPS1/2 mice at day 0 (D0), day 7 (D7), and day 14 (D14) of culture with and without treatment with pifithrin-α (PFTα). Relative gene expression compared to WT at D0. Statistics using two-tailed unpaired Student's *t* tests with Benjamini Hochberg correction for multiple comparisons. Comparisons between WT vs. HPS1 and WT vs. HPS2 at day 7 and day 14 without and with PFTα were not significantly different. Each point represents four replicate wells per time point for a total of n = 3-4 mice.

| **Gene** | **Assay ID** | **NCBI Reference Sequence** |
|---|---|---|
| *Gapdh* | Mm99999915_g1 | NM_001289726.2 |
| *Sftpc* | Mm00488144_m1 | NM_011359.2 |
| *Krt8* | Mm04209403_g1 | NM_031170.2 |
| *Cldn4* | Mm00515514_s1 | NM_009903.2 |
| *Hopx* | Mm00558630_m1 | NM_175606.3 |

**Supplemental Table 1.** qRT-PCR primer probes.
